# Mechanistic simulation identifies predictive dose-dependent biomarkers of propofol anesthesia

**DOI:** 10.64898/2026.06.10.731411

**Authors:** Anand Pathak, Scott L. Brincat, Yihan (Sophy) Xiong, Haris Organtzidis, Mason Protter, Vincent Du, Helmut H. Strey, Lilianne R. Mujica-Parodi, Earl K. Miller, Richard Granger

## Abstract

Understanding how receptor-level modulation reorganizes large-scale brain circuits remains a central challenge in neuropharmacology. Here we introduce a multiscale mechanistic model with explicit core–matrix thalamocortical architecture to examine how propofol reorganizes brainwide activity from individual receptors to systems-level circuits. Crucially, the model did not involve machine learning or statistical fitting to pre-existing anesthesia data. We use the model to simulate the multi-scale effects of propofol, which range from modulation of individual synaptic conductances to widespread alterations of neural spiking, field potentials, and coherence. Without requiring any training on task-specific data, our simulation of sensory processing in a standard auditory oddball paradigm matches independent data from anesthetized and awake behaving macaques. The same simulation, unmodified, also reproduces changes to functional connectivity in anesthetized humans, exhibiting selective attenuation of matrix thalamocortical loops relative to core loops. Most importantly, the simulation identified a previously unknown biomarker of propofol concentration: elevated residual inter-stimulus cortical activity. This dose-dependent biomarker was subsequently confirmed in empirical macaque data, where it had previously gone unnoticed. This simulation-first discovery, arising from mechanistic circuit dynamics rather than statistical comparison of clinical populations, serves as an illustrative example of a broader generative framework that can translate receptor-level modulation into circuit-scale biomarkers with potential applications across predictive neuropharmacology.

## Introduction

Pharmacological agents act at the molecular level in specific locations in the brain, yet their clinically meaningful effects encompass a broad range of complex and poorly understood consequences at the scale of distributed circuits and behaviors. Anesthesia provides a well-defined instance of this sweeping problem: the effects of anesthetic agent such as propofol on GABA-A receptors have been well characterized, but understanding the wider impact of propofol on cognition, sensory perception, and behavior requires linking its receptor-level effects with the multifaceted changes observed at higher levels. These changes include modulations of neural spiking patterns and field potentials, changes to functional connectivity, fragmentation of long-range networks, and altered thalamocortical communication(1–9). A detailed phenomenology of anesthesia-induced effects therefore involves a vast array of measurable neural and behavioral phenomena that have not been fully integrated into a single mechanistic framework.

Existing models capture fragments of propofol’s effects, but they have not yet unified the receptor, circuit, and systems levels. Connectivity analyses establish that loss of consciousness involves cortical fragmentation, but remain phenomenological: they describe what breaks down without explaining why. Mean-field approaches recover slow-wave dynamics from GABAergic kinetics (10), but such models optimize for computational tractability at the cost of the neural architecture that is crucial functionally: laminar organization and pathway-specific inhibition (5, 6). Most critically, no current model explicitly instantiates the thalamocortical core–matrix system - the circuit substrate through which propofol’s inhibitory effects are organized and propagated. Without this direct link to primary brain structural detail, the causality between molecular action and systems-level suppression remains inferential rather than mechanistic.

The cost of this explanatory gap is not merely theoretical. Clinicians titrate propofol empirically, without mechanistic guidance linking dose to circuit state to behavioral endpoint; limitations that contribute to both under- and over-sedation in practice. Most consequentially, designing next-generation agents with targeted behavioral profiles — preserving analgesia while minimizing respiratory depression, for instance — requires precisely the receptor-to-systems mapping that current models lack. A computational framework that explicitly integrates these levels would transform neuropharmacology from a largely empirical discipline into a predictive one, enabling rational drug design, biomarker-guided dosing, and principled cross-drug comparison. The framework of physiological computation (11) has previously been shown to reconcile seemingly disparate receptor, cell, and spiking effects together with field dynamics, synchrony, and behavior; we apply these methods to the domain of anesthetic perturbation. We constructed a multi-scale model incorporating multiple cortical and subcortical structures, explicitly differentiating core and matrix thalamocortical projections, laminar-specific cortical connectivity, and local and global circuit organization. Simulated propofol upregulation of GABA-A receptors is shown to cause predicted effects ranging from anterior-posterior functional connectivity reduction to gamma-band reduction to specific physiological and behavioral changes in responses to an auditory “oddball” task. Crucially, the simulation does not employ statistical training or machine learning. Instead of fitting the model to data, we built the model using the extensive literature on cell-level physiology and circuit-level anatomical organization. Data-centric modeling rarely capture known causal pathways, and for this reason they often struggle to generalize to situations not already covered in the dataset. By contrast, our mechanistic and causal approach captures how the receptor-level effects of propofol impact large-scale circuit interactions across time scales and spatial regions.

The predictions we derived from the model were validated in specific propofol experiments including functional connectivity studies in humans and auditory tasks on non-human primates. The parameters of the model were not adjusted or optimized using the experimental data. Our aim was to extract measurable predictions from bottom-up mechanistic simulations.

Our key finding is a dose-dependent physiological biomarker of propofol concentration. The likely existence of this measurable neural signal emerged from synthetic data simulated by the model, and did not involve any statistical modeling of real data. After the prediction was made, evidence for the biomarker was found in data from real propofol experiments.

Our findings therefore establish a generative framework for translating receptor-level modulation into circuit-scale biomarkers. While anesthesia is the focus here, the general modeling technique can be applied for a variety of neurophamacological applications.

## Results

### Design of the multi-scale mechanistic model

Propofol-induced anesthesia suppresses high-frequency cortical activity, leading to a behaviorally unreceptive and unresponsive state (1, 2, 12, 13). A variety of specific physiological effects have been well characterized and replicated, including changes in response amplitudes to sensory stimuli under a range of varying conditions (5–8, 14).

Propofol is an allosteric GABA-A upregulator and agonist that can directly activate GABA-A receptors and can upregulate the effect of GABA arriving at a GABA-A receptor (15–17). To study the effects of propofol on widespread brain systems, we built a multi-scale cortical-subcortical circuit model that comprises various brain structures and their anatomical connectivity at various levels, from neurons to cell assemblies to systems-level circuitry. Brain regions are modeled as cell assemblies with known anatomical connectivity at the cell, microcircuit, and multi-circuit levels. The model simulates the physiological spiking activity of individual neurons and produces simulated field potentials across multiple neurons and regions.

Propofol acts at multiple cortical and subcortical sites throughout the brain. Here, we focus on the cortex, thalamic nuclei, and specific brainstem and ascending modulatory regions. The cortical components of the model each reflect the laminar structure of cortical columns: superficial (II/III), middle (IV), and deep (V/VI) layers (see Figure 1). Each layer has its own specific input and output structures, which are largely consistent across cortical regions (18–29). Sensory inputs enter cortical columns at layer IV, relayed via core thalamic neurons (30–34). Those inputs travel up to superficial layer II/III, and then to deep layers V and VI, with each distinct input projecting topographically within a given column. Pyramidal neurons in deep layers V and VI project back to the matrix and core thalamus, respectively. Pyramidal neurons in layers II/III send cortico-cortical projections laterally to other cortical regions across the brain.

**Fig 1:**
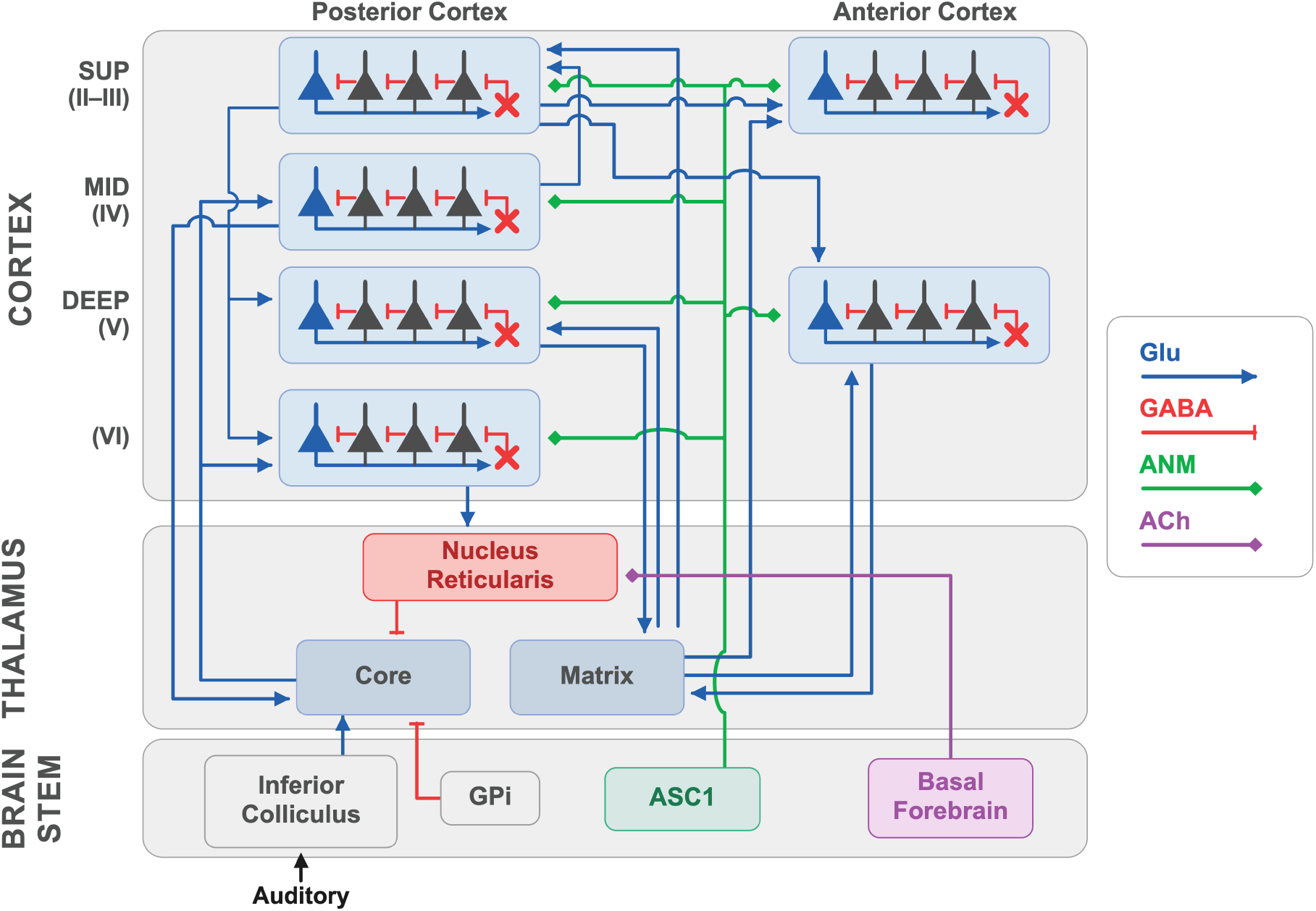
Schematic of simulated cortical and subcortical systems. Included are separate posterior and anterior cortical systems, distinct cortical layers, columnar organization, different cell types, receptor types, and connectivity, core and matrix thalamic nuclei, and the nucleus reticularis (NRt). The core thalamus predominantly receives sensory input and projects to primary sensory cortical regions; cortical feedback projects from layers V and VI to the matrix and core thalamus, respectively (see text and Methods). *GPi* : Internal Globus Pallidus, *ASC1* : Ascending input from a brainstem nucleus, *ANM* : Ascending Neuromodulator.

The model contains the two distinct types of thalamic nuclei, core and matrix, each comprised of glutamatergic neurons, as well as the reticular nucleus of thalamus (NRt) comprised of GABAergic neurons. Core thalamic neurons project predominantly to cortical layer IV, topographically, whereas matrix layers project broadly to the apical dendrites of pyramidal neurons whose cell bodies reside in layer V and in layer II/III, in a diffuse and non-topographic manner.

Core and matrix nuclei differ markedly in their spatial connectivity patterns (35, 36). Reciprocal connections between cortical layers and the core nucleus are topographically conserved across a cortical column, the nucleus reticularis, and a core thalamic nucleus. The core cortico-thalamic circuit is thus organized into modules that are anatomically and functionally modular (18–29, 37–40). Within a given vertical column, core thalamus and cortex are relatively densely connected whereas other columns, even adjacent, receive far more sparse connectivity. Primary sensory areas exhibit topographically ‘tuned’ organizations; these are retinotopic in primary visual cortex, somatotopic in touch and motor cortices, and tonotopic in primary auditory regions, such as the current simulation.

Matrix thalamic nuclei, in contrast to the core loop, project to cortical layer I, diffusely contacting synapses on apical dendrites of pyramidal cells in layer II/III and layer V (19, 36, 37, 41–43). These matrix thalamocortical synapses terminating in layer I exhibit short-term augmentation (STA): if their target neurons spike, the synapses become transiently strengthened for roughly 500 msec (44–47). (see Methods for details). Notably, these transient changes enable these synapses to participate in temporal encoding of stimuli (see (37)).

### Simulation independently replicates distinct brain response features observed in macaque as well as human propofol experiments

The simulation was presented with a set of auditory stimuli under simulated ‘normal’ and ‘propofol anesthetized’ conditions. These stimuli corresponded with the stimuli presented to awake and propofol-anesthetized macaques (14). Each stimulus consisted of five rapid tones, either all the same (A-A-A-A-A) or one different (A-A-A-A-B), i.e., the ‘oddball’ (see Figure 2a-d).

**Fig 2:**
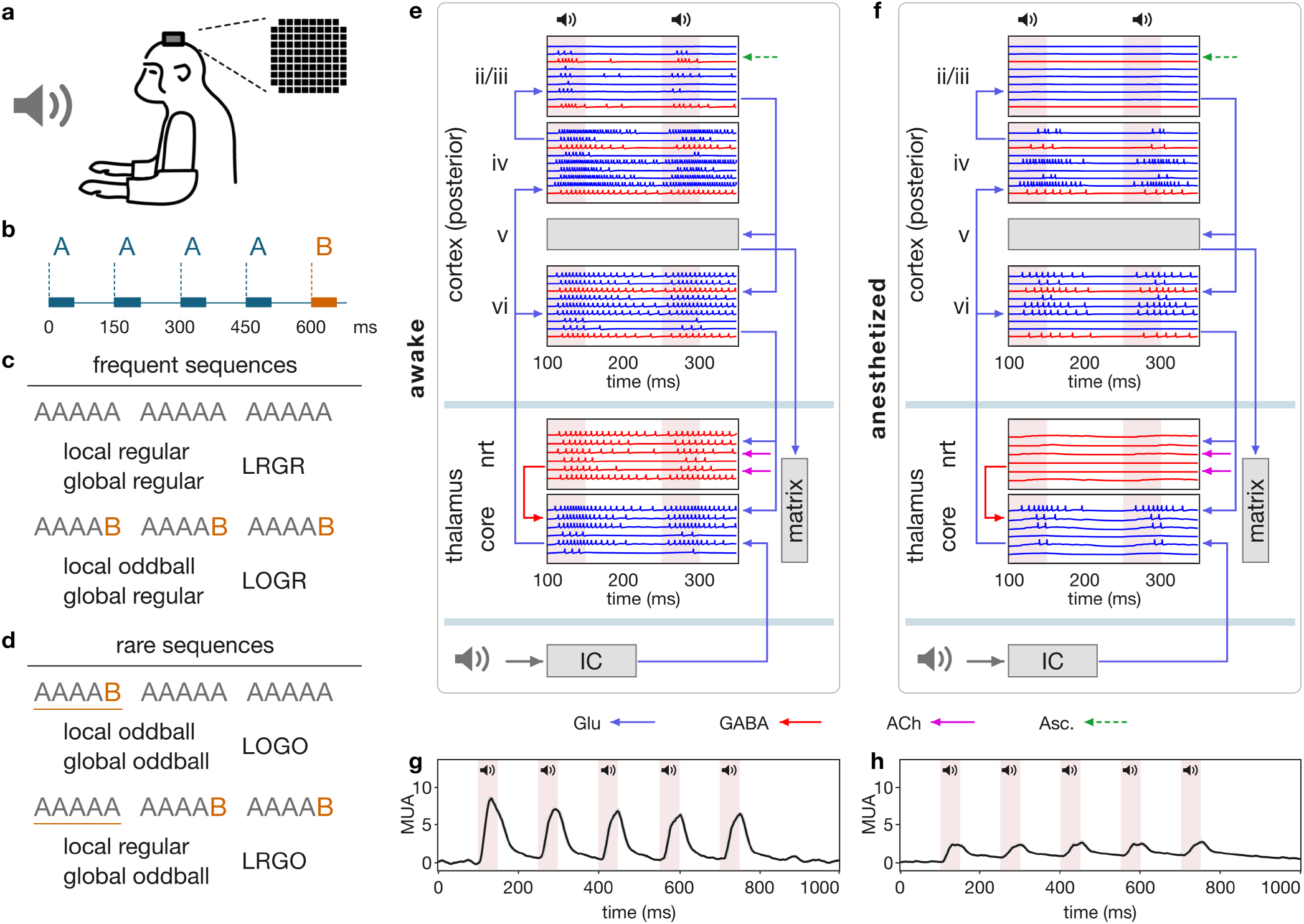
Bottom up modeling of alteration in the brain activity, emerging from the receptor level modulation. Propofol modulated increase in amplitude and time courses for GABA-A receptors inhibitory postsynaptic currents (IPSCs; see Supplementary Figure S4) causes broad changes in response of multiple brain areas. (a-d) Experimental paradigm presented to the simulation and to macaques. “Oddball” tones (B) were systematically interspersed into repeated regular tones (A) in temporal patterns that could be varied to make the oddball locally (within a sequence) or globally (across multiple sequences) unexpected. (e-f) Spiking activity (and thus power in the gamma range) in core loop (predominantly posterior) cortical and thalamic areas (see Figure 4 a) is substantially altered by propofol: spiking is reduced throughout superficial cortical layers (ii-iii), in the thalamic nucleus reticularis (nrt), and in core thalamic nuclei. Bottom panels show simulated field potentials in awake (g) and propofol (h) conditions measured from core (posterior) cortical regions in response to a single oddball auditory trial in the local regular global regular (LRGR) condition, i.e., where AAAAA occurs frequently. Shaded pink regions indicate duration of auditory stimulus. See text for description and discussion.

The four possible sequence patterns are: local oddball, global regular (LOGR); local regular, global regular (LRGR); local regular, global oddball (LRGO); local oddball, global oddball (LOGO). Intuitively, these can be understood in terms of repetition versus rarity. Global oddball local oddball (LOGO) contains occasional AAAAB trials in a block of mostly AAAAA trials; global oddball local regular (LRGO) have occasional AAAAA trials in blocks of mostly AAAAB trials. Note that in LOGO, the overall occurrence of the B sound is very rare, whereas in LRGO, B is far more common – the rarity only arises from its occasional absence (in an AAAAA trial). (In LRGO trials, the ‘oddball’ is considered to be not the oft-heard B, but the trials of AAAAA, that drop the B.) The global regular sequences are local regular global regular (LRGR) and local oddball global regular (LOGR); the former contain no B sounds at all, just repeated sequences of five As. LOGR contains regularly repeating sequences of AAAAB; thus, over repeated blocks, the B is no longer particularly rare. Thus, the only trials with truly rare B occurrences across blocks are “LOGO” trials, i.e., those that contain occasional AAAAB interspersed among mostly AAAAA trials.

Figure 2e-g illustrates the model simulation of a single LRGR trial, showing the effect of increasing GABA-A inhibitory postsynaptic currents (IPSCs) (Supplementary Figure S4). It can be seen that spatiotemporal firing patterns are substantially reduced by propofol throughout the core corticothalamic circuit (Figure 2e-f). Propofol is known to reduce gamma spiking activity, leading to the use of gamma activity as a marker of anesthesia (6, 14). Our simulations also demonstrated this phenomenon in both posterior and anterior cortices (Supplementary Figures S6 and S7). Moreover, population activity is correspondingly reduced, as seen in simulated physiological field responses in the awake (Figure 2g) and propofol anesthetized (Figure 2h)) states. Propofol’s enhancement of GABA-A responses has an net suppressive effect on cortex, but in other brain regions a disinhibitory effect can dominate as a result of increased suppression of inhibitory neurons. For instance, nucleus reticularis becomes largely quiescent under propofol, due to increased GABA-A input from ascending modulatory systems (see Fig 1, ASC2). In general, the interplay among excitation, inhibition and disinhibition mean that a centrally active compound can have different effects in different regions, as opposed to a uniform excitatory or inhibitory effect. Our simulations capture an example of this kind of interplay and its consequences at multiple scales.

We provide a preview of the major findings in Figure 3. No data-driven statistical fitting or machine learning was used to tune parameters, yet the model was capable of matching empirical findings. Shown are the simulated (left) and empirical macaque (right) physiological responses to sequences of five tones in various arrangement patterns (see Figure 2a-d). It can be seen in Figure 3 that these LOGO trials elicit the largest responses to the (rare) B sound (see the purple (LOGO) responses marked as 2, 4, 6 in the empirical portion of Figure 3). (The next most prominent responses are LOGR (red), which repeatedly play the oddball B after repeated As. The LRGO (blue) trials are those where the normal AAAAB is replaced by a (supposedly oddball) AAAAA; these can be seen to elicit no notable response, which intuitively is because the final (fifth) A is locally not surprising, and is globally not at all rare.)

**Fig 3:**
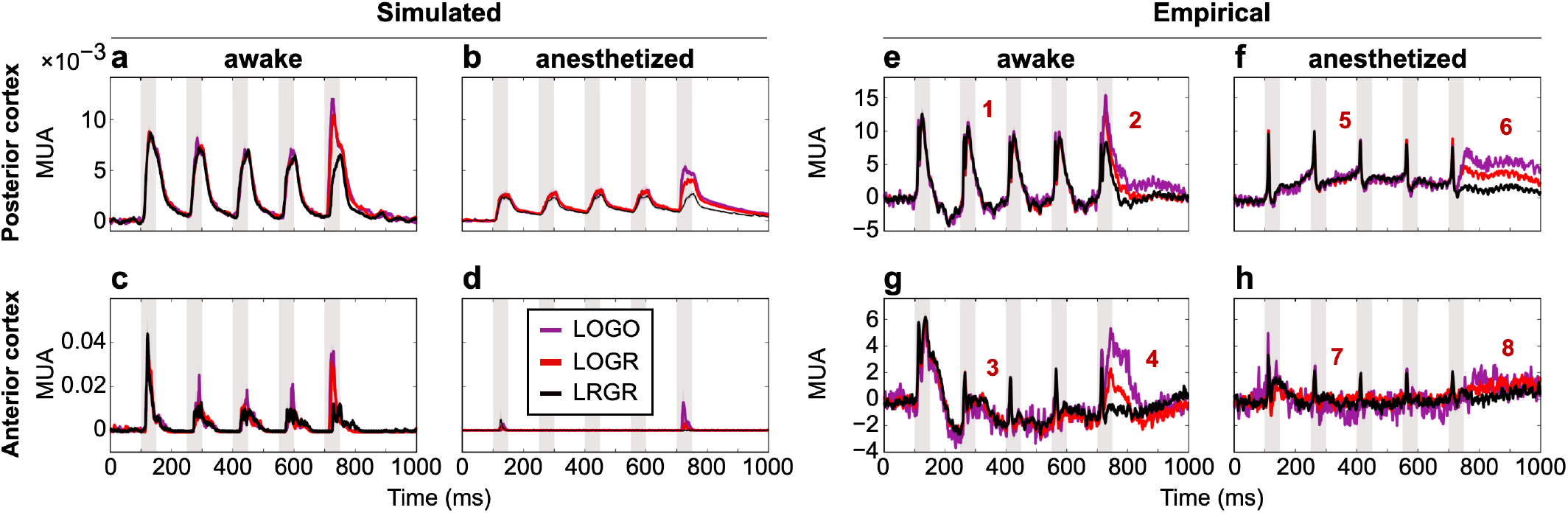
The model exhibits distinct features observed in empirically measured brain responses to auditory stimuli during awake and anesthetized states. Side by side comparison of simulated responses to tone sequences (a-d), and the corresponding responses measured in macaques (e-h), in both posterior and anterior cortical regions, under control awake and anesthetized conditions. Shaded pink regions indicate the duration of each auditory stimulus. The model was presented with the same auditory tone sequence stimuli that were presented to the macaques (see Fig 2), resulting in a mechanistic simulation of the physiological responsed that did not require any data-driven statistical fitting or machine learning. The factors that give rise to the simulated responses in key epochs (marked with numerals here) are described in detail in the text. These factors serve as testable hypotheses about of the causes of the experimentally observed responses in macaques. (red, LOGR; gray, LRGR; purple, LOGO; see text for abbreviations).

Here we provide purely mechanistic explanations of how each of the prominent physiological effects occurs in the simulation. These explanations serve as testable hypotheses of how the corresponding effects may arise in the empirical macaque physiological measures. The eight effects (as indicated in Figure 3e-h) are (1) Attenuation of posterior cortical responses to stimulus repetition in the awake state; (2) Rebound responses to oddball stimuli in posterior cortex; (3) Strong attenuation of anterior cortical response to stimulus repetition in the awake state; (4) Rebound responses to oddball stimuli in anterior cortex in the awake state; (5) Absence of posterior cortical response to repeated stimuli in the anesthetized state; (6) Rebound response to oddball stimuli in posterior cortex in the anesthetized state; (7) Suppression of anterior cortical responses to repeated stimuli in the anesthetized state; (8) Modest rebound response to oddball stimuli in anterior cortex in the anesthetized state.

### Attenuation of posterior cortical responses to stimulus repetition in the awake state

This reduction of response to repeated stimuli in posterior sensory cortex, indicated in Figure 3e (epoch 1), occurs almost entirely due to the successive build up of inhibition within the core nucleus by the nucleus reticularis projections, which in turn inhibits thalamic responses to inputs. The auditory stimulus activates core thalamic nucleus through inferior colliculus (See Figure 1 and Figure 2e-f), which then activates deep layer VI of posterior cortex. Layer VI pyramidal neurons then activates the nucleus reticularis in the thalamus that project back to core nucleus. Nucleus reticularis neurons generate inhibitory post synaptic potentials with a prolonged time course (48, 49) that is extended further as a result of summation of successive stimuli that arrive within a small time window (in this case 150 ms, see Supplementary Figure S1)

### Rebound responses to oddball stimuli in posterior cortex

The posterior cortical rebound response to oddball tones(Figure 3e, epoch 2), arises in large part due to short-term potentiation within the inferior colliculus. Inferior colliculus adapts to the statistics of auditory stimuli, responding differentially to repeated versus rare stimuli (50). Oddballs (both global and, to a lesser extent, local) activate distinct posterior cortical modules, such that thalamic response to oddball tones are not subject to the built-up inhibition from nucleus reticularis, in contrast to the thalamic response to repeated tones, as described above (see Supplementary Figure S2)

### Strong attenuation of anterior cortical response to stimulus repetition in the awake state

The rapid suppression of the anterior cortical response to repeated stimuli (Figure 3g, epoch 3), arises from an interaction among three key factors: i) short-term augmentation of thalamocortical synapses (44–47), ii) overlap between thalamocortical and corticocortical anterior targets (19, 36, 37, 41–43); and iii) lateral inhibition within anterior cortical columns (51–55). The anterior cortical cells that receive convergent input from both thalamus and posterior cortex are far more responsive than cells that receive only one or the other. These highly responsive cells laterally in turn boost inhibition on their neighbors (11, 56) (see Supplementary Figures S3 and S5.)

### Rebound responses to oddball stimuli in anterior cortex in the awake state

This rebound response (Figure 3g, epoch 4), is closely related to the suppression of the posterior cortical response to repeated tones described above. Oddball stimuli (both global and, to a lesser extent, local) activate distinct posterior cortical modules, which then selectively converge onto anterior cortex alongside diffuse thalamic input. The anterior cortical cells that respond to the oddball tone are distinct from those that respond to repeated tones so the corresponding anterior cortical cells do not experience a substantial strengthening of thalamocortical synapses, or a boost in local lateral inhibition

### Absence of posterior cortical response to repeated stimuli in the anesthetized state

Propofol-induced changes to posterior cortical responses (Figure 3f, epoch 5) are caused by increases to GABA-A responses in both cortex and thalamus (15–17). This effect can also be seen in Figure 2

### Rebound response to oddball stimuli in posterior cortex in the anesthetized state

Propofol-induced cortical rebounds (Figure 3f, epoch 6) arise predominantly because cells in the inferior colliculus (I.C.) learn the statistics of repeated stimuli and suppress their responses to them. The same mechanism also causes the rebound response to oddball tones in posterior cortex (Figure 3e). Oddball stimuli arrive at cells in inferior colliculus whose responses have not been supressed: these responses are strong enough to transiently overcome cortical inhibition

### Suppression of anterior cortical responses to repeated stimuli in the anesthetized state

Blanket inhibition of most layer II/III anterior cortical activity (Figure 3h, epoch 8) is induced by propofol. Layer II/III contains inhibitory interneurons that enable strong feedback inhibition as well as lateral inhibition among pyramidal neurons (51–55) and propofol-induced GABA-A agonism in layer II/III therefore causes strong suppression of any activity

### Modest rebound response to oddball stimuli in anterior cortex in the anesthetized state

Highly suppressed anterior cortical oddball rebound (Figure 3h, epoch 8) is again enabled due to topographically disinhibited inferior colliculus cells,(Figure 3f, epoch 6)

Taken together, these findings suggest that anesthesia has a differential effect on matrix versus core thalamocortical activity. We simulated functional connectivity corresponding to the method used by (7). Figure 4b shows that the functional connectivity in the matrix circuit is highly disrupted due to propofol induced anesthesia, while that in core circuit is weakly affected in comparison. Figure 4c demonstrates a validation of these predicted effects on core and matrix loops: the same qualitative effects were reported in an independent study of propofol effects in humans (7). Our simulation predicted the incremental dose dependent suppression of matrix loop FC upto a specific dose factor (see Figure 4d), beyond which there is a rapid disruption of matrix loop functional connectivity.

**Fig 4:**
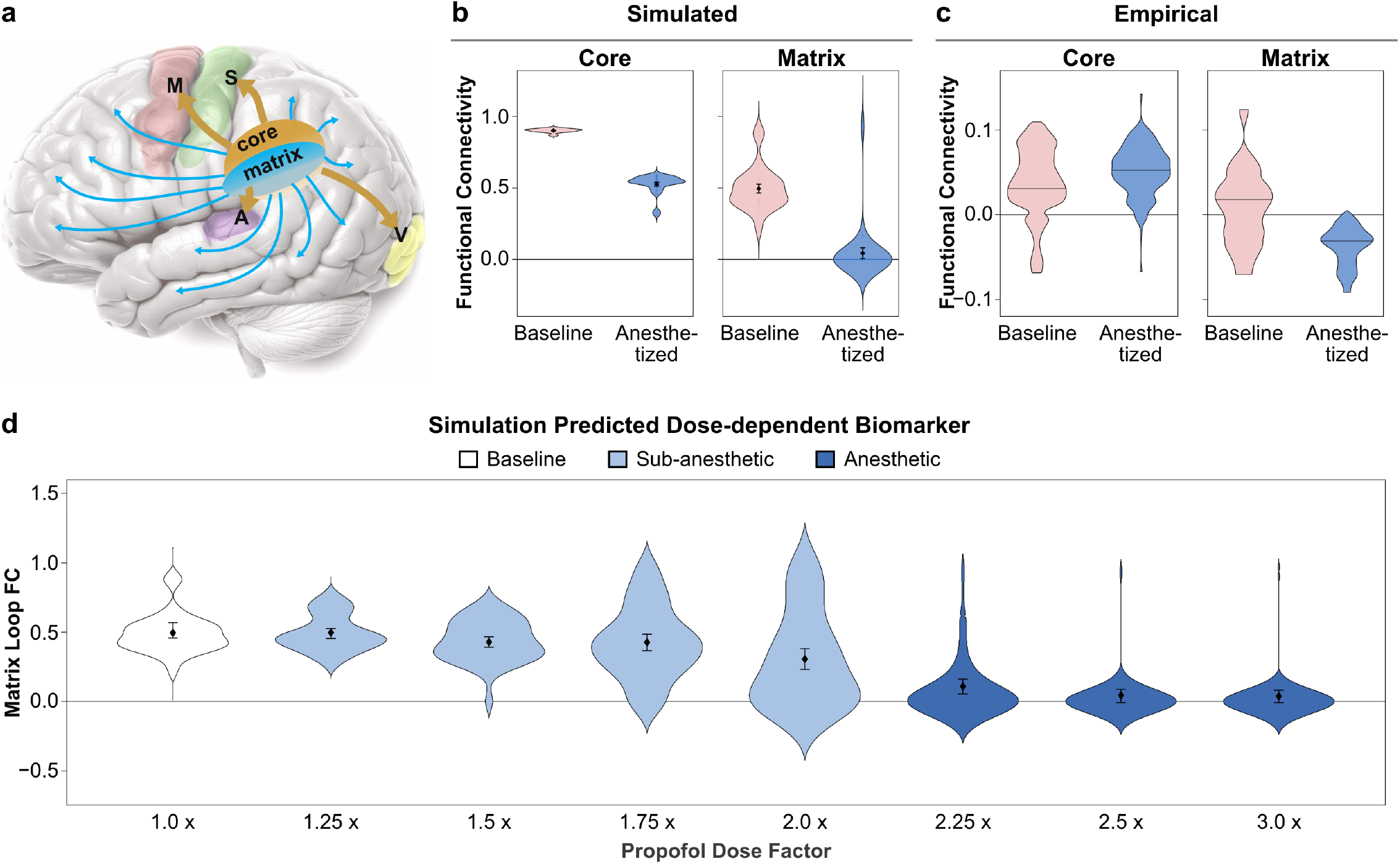
Simulations predict propofol mediated disruption of matrix loop functional connectivity, observed independently in human fMRI. (a) Thalamic core nuclei project topographically to primary sensorimotor cortical areas: motor (M), somatosensory (S), auditory (A), visual (V); by contrast, matrix thalamic nuclei project broadly and diffusely to all other cortical areas. (b-c) Side by side comparison of simulated and empirically measured functional connectivity (FC) of core vs. matrix thalamocortical activity patterns. The empirical results on human subjects shown here are taken from reference (7) (see Figure 6 in (7)). Functional connectivity in Core and matrix is differentially affected by propofol anesthesia; core is little affected whereas matrix FC gets strongly reduced. (d) Simulation predicted incremental disruption of marix loop FC with increasing dose of propofol. Dose here is expressed in terms of dose factor i.e. the factor by which the IPSC amplitude and timecourses of GABA-A receptors are increased at a given propofol concentration.

### Residual activity emerged as a direct biomarker of anesthetic dose

The simulation results presented so far mirror the strong brain-wide effects of anesthetic doses of propofol. We now show that our model can also simulate sub-anesthetic doses accurately. Propofol’s modulatory effect on GABA-A receptors varies monotonically with concentration, making it tractable to simulate the effects of different doses.These simulations allowed us to identify a dose dependent biomarker: residual cortical activity in the post-stimulus period. This biomarker was seubsequently found in the empirical data. We validated our biomarker at a low dose with the empirical recordings done in oddball trials 40-50 minutes after the cessation of propofol infusion. After this time interval, upon the cessation of drug infusion, propofol concentration in the brain is pharmacokinetically estimated to be roughly 50% of the initial anesthetic concentration (57). Propofol concentration directly affects synaptic conductance as assessed by measures of the amplitude and decay-time of inhibitory postsynaptic currents (IPSCs, see Supplementary Figure S4) (15–17). We simulate the brain-wide effects of alterations to these physiological parameters, spanning from awake, to low dose (sub-anesthetic), to high dose (anesthetic) concentrations.

Figure 5 shows the measureable neural signals that reflect the dose dependent effect of propofol: the amplitude of the responses to oddball stimuli in posterior cortex, and more prominently, elevated residual activity underlying those responses, as measured between pairs of successive stimuli (see the rectangles in Figure 5 a and b). Figure 5 a and c demonstrate that the model replicates the dose dependent behavior, i.e. reduced cortical response and increased residual post stimulus activity between consecutive stimuli. Figure 5 c-d show that residual activity acts as a direct biomarker of propofol dose.

**Fig 5:**
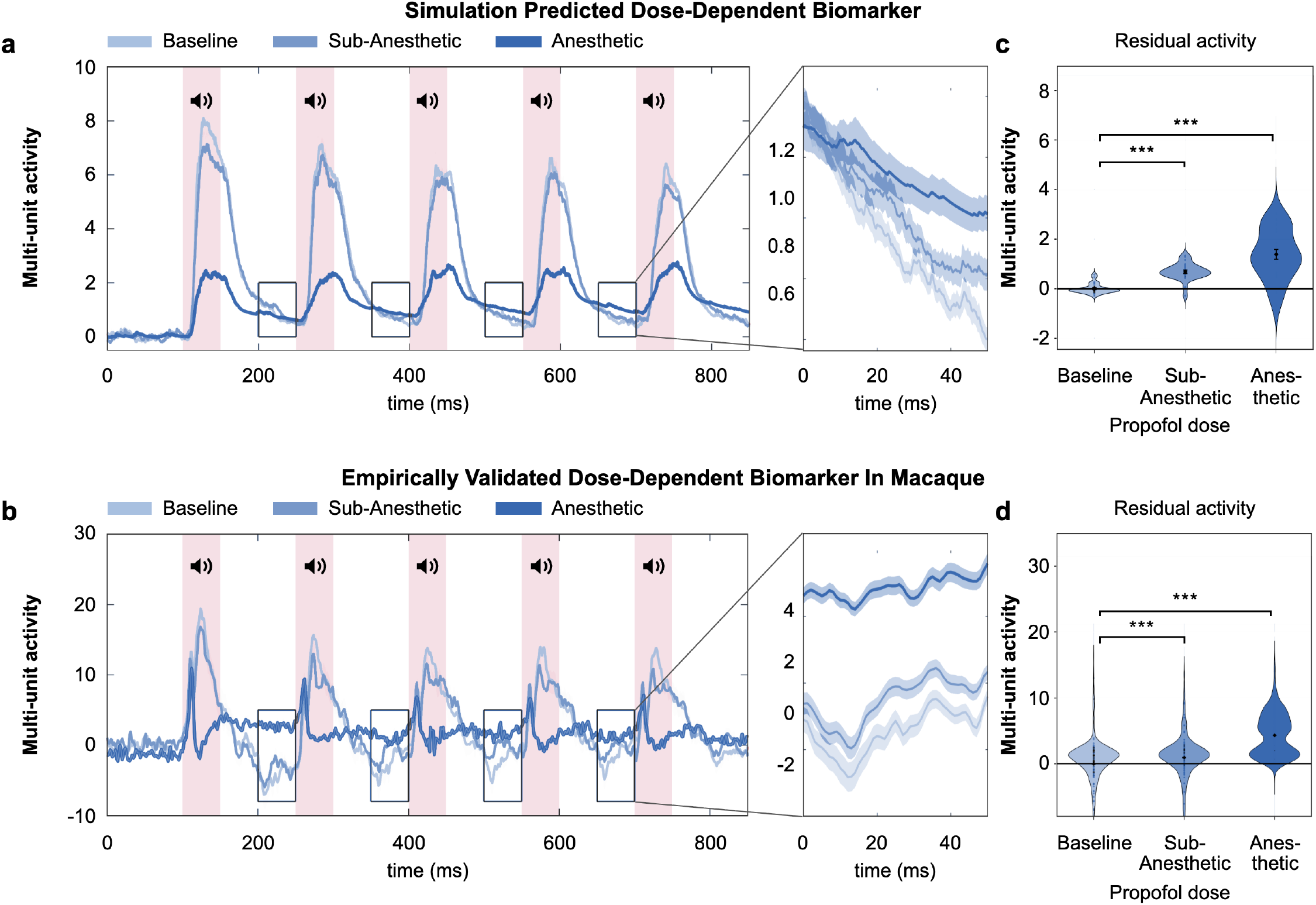
Identification of dose-dependent biomarker in propofol simulations and subsequent validation with empirical results. (a-b) Simulated and empirical averaged responses to single auditory five-tone sequence (AAAAA). (The tones are on for the fifty milliseconds shown in shaded pink vertical bars, and then off between these.) Responses are shown for unanesthetized (baseline, light blue), fully anesthetized (dark blue), and sub-anesthetic concentration (blue) during recovery (see text). The recovery time intervals are marked by small black boxes and magnified in the adjacent panels. The magnified “residual” portions of responses from the small black boxes shown in panels a and b, indicate the residual background activity that occurs between auditory-driven responses during the 50 msec immediately before the subsequent tone in the sequence, for simulated and empirical responses. It can be seen that the awake (baseline) residuals are lowest, fully anesthetized residuals are highest, and the sub-anesthetic dose is intermediate. (c-d) Net residual activity from the average awake baseline; awake (light blue) is at baseline, sub-anesthetic dose (blue) is higher, and full anesthetic dose is highest (dark blue), in both simulation (c) and empirical (d) measures. The shift in the residual activities from the base line are statistically significant (p<0.001) as measured by two-sample KS test. The simulated residual responses appear to be predictive of the empirical residual responses. The shift in the residual activity in different doses are statistically significant, which qualifies it as a reliable biomarker for anesthetic dose. See Supplementary Figure S8 for simulation results of incremental shifts in residual activities with increasing doses of propofol.

The simulated responses arise relatively straightforwardly from *bottom-up simulations that were not trained on any macaque data* (as in previous related work (11)). In the simulation, each effect can be mechanistically understood without reference to data fitting. The resulting mechanistic explanatory accounts of these effects constitute explicit hypotheses for how the corresponding empirical macaque effects arise. We further performed a strong perturbative test for the robustness of our model by simulating an ensemble of models, where in each run we simultaneously perturbed connection parameters by 20% to 50% from our original model. Even with such strong perturbations, while the circuit topology remaining intact, we obtained all the qualitative results presented above (see Supplementary Figures S9-S11)

## Discussion

We demonstrate a mechanistic circuit model that is capable of simulating brain activity across multiple scales: physiologically realistic spiking cells are linked together in anatomically grounded micro-assemblies that form a system-level circuit capable of simulating brain-wide activity. The circuit model is used to simulate the neural basis of sensory perception in a passive acoustic task. The model was constructed from accurate physiological and anatomical data corresponding to specific cells as arranged in specific brain structures, each of which performs a basic functional computation that emerges from its biological activity alone, i.e., physiological computation. These basic computations themselves combine and interact to give rise to complex behavior across multiple brain areas. The neural responses simulated by the model matched independent experimental data from the literature on human subjects and on macaques under propofol anesthesia (7, 14). These simulated responses map directly onto experimentally measurable quantities and include detailed spiking behavior, local field potentials, cortical functional connectivity, and responses elicited by sensory stimuli. Crucially, this correspondence did not arise from direct data-driven fitting: no machine learning or statistical parameter estimation was employed to calibrate the model. As a result, the predictions generated by a single, unmodified model cover both nonhuman primates and humans, and apply to different types of measurements, including electrophysiology (local field potentials) and functional connectivity.

Anesthetized subjects can selectively respond to auditory cues and can register physiological markers of “surprise” at unexpected sounds in otherwise-predictable sequences, as in the oddball task used here. Measures from anterior and posterior cortical areas of macaque brains exhibited particular findings in awake and anesthetized states; these were closely matched by the simulation. Notably, the simulations reveal a marked difference between anterior and posterior cortical effects (e.g., top vs. bottom of Figure 3). These simulated anterior-posterior differences arise largely from the anatomical distinction between matrix and core thalamocortical projections. In particular, matrix thalamus exhibits diffuse projections to anterior cortex whereas core thalamus projects predominantly topographically to specific cortical sensory regions (Figure 4a). In the simulations, propofol had a very strong suppressive effect on matrix thalamocortical responses; the analogous effect on core thalamocortical activity was much weaker. The simulated rebound responses to oddball stimuli, in both awake and anesthetized states, were far more attenuated in anterior cortical activity than in posterior responses (compare points 2 and 6 to points 4 and 8 in Figure 3). In addition to these predictions, which were verified in macaques, the same model, with no changes, also predicted a clear distinction between core and matrix circuit loops in functional connectivity that was borne out in data from propofol-anesthetized humans (Figure 4b-c), even though the model was not constructed using this type of data.

Most importantly, the model enabled us to identify a novel biomarker that distinguishes among propofol doses in responses to repeated auditory tones (Figure 5a). Evidence for this candidate biomarker, which had eluded previous analyses of the data, was successfully found in the empirical data (Figure 5b). Most drug effect biomarkers in the literature are derived from statistical comparisons among different populations, as is often done during clinical trials for pharmaceutical drugs. In our case a biomarker was discovered without large-scale clinical trials; instead, the biomarker emerged from the analysis of mechanistic simulations, prior to the examination of empirical data.

These diverse findings, which include the de novo identification of a novel dose-dependent predictive biomarker, arose from mechanistic simulation of physiological operations occurring in neural regions connected in an anatomically realistic circuit. This approach, which we term “physiological computation,” stands in contrast to popular approaches in artificial and biological neural network systems, such as statistical training, gradient descent and reinforcement learning. The present model is not designed or trained with a goal of capturing statistical patterns in large datasets, an approach that dominates machine learning and AI. Instead of fitting a target dataset, the model simulates realistic physiological activity; its behavior emerges solely from its grounding in the detailed physiology of neurons and synapses, and their anatomical connectivity. The model receives simulated auditory stimuli and then generates responses that begin in the sensory periphery, propagating through the circuit via well-established anatomical projections. The correspondences between the simulation’s responses and actual empirical measures provide a demonstration that realistic physiological spiking, embedded in anatomically accurate circuitry, can give rise to the physiological responses that brains produce.

Our aim here was to develop a mechanistic understanding of how centrally-active compounds affect brain circuitry. Receptor-specific drugs act locally at specific sites in the brain, but there is no purely data-driven way to extrapolate from local observations to their large-scale consequences. Methods that take training data and identify how to statistically mimic it are highly underconstrained: hundreds of different data models all are capable of fitting the data and yet the approaches typically select just a few. Analysis of how receptor modulation affects multiple scales of circuit activity may require detailed mechanistic modeling of brain structures, each composed of distinct cells arranged in heterogeneous anatomical patterns. Many modeling approaches smooth away this mechanistic heterogeneity through averaging, mean field approximations, and artificial neural networks (9, 58). Some other models share our goal of adopting careful anatomical and physiological constraints (10, 59), as opposed to using simplifying abstractions such as mean fields, artificial neural networks, or abstract dynamical systems. However, such approaches leave out many crucial neurobiological details, and are therefore unlikely to capture the counterintuitive phenomena that often arise in real data. Though many modeling approaches map manipulations of a neural model onto their measurable outcomes, it is still relatively rare for models to span scales from receptor dynamics to large-scale anatomical circuit activity.

Mechanistic modeling of the kind we demonstrate here is crucial for basic science, but our approach also has important translational applications. Computational models have largely had a limited ability to assist researchers focused on real-world applications such as pharmaceutical trials. This may be a contributing factor to the longstanding disconnect between data-fitting approaches to neural activity, on one hand, and predictive analyses of pharmacological drug effects in the brain on the other.

This in turn may contribute to the extremely high (70-80 %) failure rate of Phase II drug trials after successful Phase I trials (60).

The propofol simulations reported here illustrate the clinical applicability of our general modeling approach. In the clinic, anesthesia is typically administered as a bolus, leading to the rapid onset of unconsciousness. By contrast, the macaque studies cited here use slower infusion enabling study of intermediate states of induction and recovery. Similar slow-infusion studies have been carried out in humans; such studies report a substantial shift in alpha power to anterior areas and its reduction in posterior regions, simultaneously with onset of unconsciousness (1, 61). These and other notable electrophysiological measures are topics of ongoing and future studies.

Hundreds of millions of people per year receive anesthesia, rendering patients unconscious yet carefully maintaining heart rate, blood pressure, respiration, and other crucial somatic functions. Such treatments require the expert attention of specialists who monitor biological function to ensure patients’ health, but identification and quantification of accurate dose-dependency of anesthetic effects remains a topic of active study (14, 62). The present findings of specific mechanistic dose-dependent biomarkers of propofol anesthesia may help advance our understanding of these patient-crucial anesthetic characteristics, as well as providing a platform for testing of novel centrally active compounds.

## Materials and Methods

### Detailed Model Description

#### Spiking neurons and synaptic connections

##### Neuronal Dynamics

Each neuron is modeled using ionic conductance based point neuron models following the Hodgkin-Huxley formalism (63). For simplicity, in this model we consider only voltage gated Sodium and Potassium channels. The membrane potential is described by:

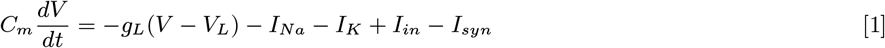

where *I*_*in*_ is the input current, which can either be external stimulus or background activity, and *I*_*syn*_ is the synaptic input.

Sodium current 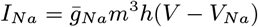and potassium current 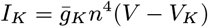are driven by gating variables {*m, h, n*} whose dynamics are governed by the general form :

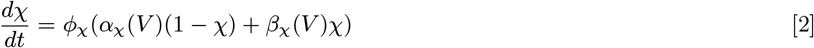

Here, *χ* is the gating variable and *ϕ*_*χ*_ is the temperature related effect on the time scale.

##### Synapse

Synaptic current in Eq. 1 for a neuron *i* is the sum of synaptic inputs from all incoming axons *{j}* viz. 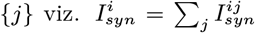.

Synaptic current from neuron *j* to neuron *i* is given by

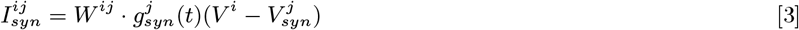

where *W* ^*ij*^ is the synaptic weight from neuron *j* to *i*, a quantity which estimates density of dendritic spines and is modified by Long Term Potentiation (LTP) and Long Term Depression. The synaptic conductance 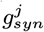 is a function of spiking activity of the presynaptic neuron, and therefore is one of the state variables for each neuron whose dynamics is described by damped oscillator equation:

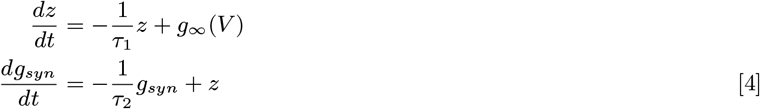

which is driven by impulse force *g*_*∞*_(*V*) defined as follows (64)

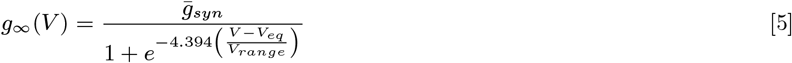

##### Cortical areas

Superficial layers of cortical regions in this model (cortical blocks) comprise large number of local feedback lateral-inhibition circuits (L-FLIC). For detailed explanation see reference (11). Each L-FLIC circuit comprises 5 excitatory neurons sending synapses to one feedback inhibitory neuron, while the inhibitory neuron sends synapses back to the 5 excitatory neurons. The L-FLIC circuits are connected to each other through synapses between the excitatory neurons. We have set the connection density such that each excitatory neuron is connected to 1 excitatory neuron from the same cortical block. Within each cortical block, we have 1 *feed-forward* inhibitory neuron that receives input from excitatory neurons in other cortical regions and forms inhibitory synapses onto excitatory neurons within the same cortical block. These feed-forward neurons also receive modulatory inputs from various *Brainstem* nuclei forming the Ascending Reticular Activating System (65). The rhythmic bursting of feed-forward inhibitory neurons causes the rhythmic bursting of the excitatory neurons and which in turn sets the dominant rhythm in the cortical region.

###### There are 5 layers in each cortical column. In the posterior cortex, we model them as follows

###### Superficial Layer II-III

This layer is identical to the cortical block developed in our previous model of the corticostriatal circuit (11). It comprises 10 LFLIC circuits i.e. 50 pyramidal neurons and, 10 GABAergic feedback interneurons and 1 GABAergic feedforward interneuron.

###### Middle Layer IV

The middle layer is similar to the superficial layer in its underlying structure. It comprises 5 LFLIC circuits (25 pyramidal neurons and 5 GABAergic feedback interneurons). However, it has marginally higher local connectivity among the pyramidal neurons (*density* = 3%), compared to extremely sparse local connectivity seen in the superficial layer (*density* = 1%). The feedback inhibition is significantly weaker than in the superficial layer.

###### Deep Layer V

Layer V is similar in structure to Layer IV (5 LFLIC circuits) with higher local connectivity between pyramidal neurons (*density* = 5%). Feedback inhibition is significantly weaker than in the superficial layer.

###### Deep Layer VI

Similar in structure to Layer V with extremely low feedback inhibition.

These cortical layers, connected together, form a single module within the posterior cortex that corresponds to a single sensory input (the frequency of the auditory stimulus in this case).

#### Thalamic nuclei

Thalamus comprises three primary structures (see the model details in the Results section), Core, Matrix and the Nucleus Reticularis (NRt). Core and Matrix consist of Glutamatergic neurons while NRt consists of GABAergic neurons. All three also differ in their input and output projections.

##### Core Nucleus

Core is modeled as a sparsely connected network of 25 Glutamatergic neurons (connection density = 2%).

##### Matrix Nucleus

Modeled same as core with 25 glutamatergic neurons.

##### Nucleus Reticularis

Comprises 25 GABAergic neurons. They have long synaptic decay *τ*_*inhib*_ = 250*ms* compared to other GABAergic neurons who have *τ*_*inhib*_ = 70*ms*

All the cortical layers of the posterior cortex and three thalamic nuclei have modular structure, such that they are anatomically and functionally bundled (37). Therefore we have two separate modules for all the above structures, one corresponding to each auditory stimulus frequency. We have two blocks for each cortical layer in posterior cortex and for core, matrix and NRt. For anterior cortex we have only one block of superficial layer II-III.

#### Brainstem regions and GPi

##### Inferior Colliculus

Modeled as a simple source of input what projects to core. We modeled it as a pulse current source whose amplitude is monotonically decreasing function of stimulus frequency. This simulates the adaptation in IC described above in Results section (50).

##### ASC1

This is a modulatory input to superficial cortical layers. The simulated ascending system (ASC1) is a simplification of reticular ascending systems such as the cholinergic basal forebrain and noradrenergic locus coeruleus; it produces rhythmic activity using a neural mass model that feeds modulatory input to cortex, driving a beta rhythm in the cortex(66). The ASC1 simulation thus functions as a central pattern generator. It has been modeled using a simple ‘Next Generation’ neural mass model (67, 68). It has been modeled using same equations as done in our earlier model in reference (11)

##### ASC2

This is another simplified brain stem region that gives tonic excitatory input to NRt.

##### GPi

We modeled a simplified GPi mass that gives a steady inhibitory input to core thalamus in the model.

#### Simulating Propofol effect

We simulate the effect of propofol through a single parameter *P*_*gsyn*_ that changes the amplitude and the decay rate of GABAergic IPSCs throughout the circuit for every GABA-A synapse. For awake state the parameter by default is 1 and for anesthetized state *P*_*gsyn*_ = 2.5. For sub anesthetic effect at propofol concentration in brain at *≈* 50% of anesthetic dose, *P*_*gsyn*_ = 1.5. Both ASC2 and GPi get affected differently by propofol. ASC2 activity is suppressed (due to presumed GABAergic neurons within ASC2) which shuts down the activity of NRt. GPi input gets strengthened since they are comprised of GABAergic neurons. Thus, even though they are not modelled explicitly as neural assemblies the effect of propofol is accounted.

##### Experimental task design

The auditory oddball task and the corresponding neurophysiological recordings in adult rhesus macaques, which we reanalyzed and used here for validation, are described in detail in ref. (14) and summarized here. The animals were cared for in accordance with National Institutes of Health guidelines and the policies of the Massachusetts Institute of Technology Committee on Animal Care. Two monkeys passively listened to sequences of five pure tones, adapted from the local–global paradigm of (69). Each tone lasted 50 ms and was followed by 100 ms of silence before the next tone, and successive sequences (trials) were separated by 500 ms. Tones were either A (800 Hz) or B (1600 Hz). Each block began with 20 presentations of a frequent sequence (AAAAA, BBBBB, AAAAB or BBBBA) to establish the global regularity. These were followed by a random mixture of 80 frequent and 20 rare sequences; the rare sequence shared the first four tones but differed in the fifth (e.g., AAAAA within an AAAAB block). Crossing local deviance (fifth tone different or not) with global rarity gives the four trial types used here: LRGR, LOGR, LRGO and LOGO. Spiking and multi-unit activity were recorded with Utah arrays and laminar probes in auditory parabelt cortex (area Tpt) and frontal cortex (area FEF), which serve as the posterior and anterior cortical regions in our comparisons. Propofol was infused intravenously at an induction dose for about 20 minutes and then at a lower maintenance dose for about 40 minutes. Loss of consciousness was defined as behavioral unresponsiveness and spontaneous eye closure. Recordings were obtained in the awake state, during propofol-induced unconsciousness, and during recovery after the infusion ended, from which the sub-anesthetic epoch (40–50 minutes after cessation) was taken.

##### Simulation Protocol

We presented the simulation with the same tone sequences described above. Each trial comprises 5 successive tones of same frequency (LRGR type, see above) or 4 successive tones of same frequency followed by 5th tone of a different frequency (LOGR and LOGO type). Each tone is on for 50 ms followed by 100 ms of silence before the next tone. We then measure the simulated output for a region by averaging the postsynaptic currents within that region for all the underlying neurons. Each tone frequency is mapped onto a separate cortico-thalamic module, as mentioned above. For instance tone A activates IC-A, that feeds into Core Thalamus-A, which further activate the posterior cortical layers of module A, and there is a similar circuit activity for tone B which activates posterior cortical module B. Thus by setting the activity pulses for both the IC modules, we can simulate all different types of oddball trials.

##### Functional Connectivity

We calculated the functional connectivity shown in Figure 4 using Pearson’s correlation coefficient of the simulated multiunit activity for posterior cortex, anterior cortex, core thalamus and matrix thalamus. For core connectivity we measure correlation between posterior cortex and core while for matrix connectivity we measure correlations between anterior cortex and matrix.

##### Residual Activity

The residual activity between two auditory stimuli was calculated as a dose dependent biomarker for both simulated and empirical MUAs. For empirical we followed the same baselining and normalization process as was done in (14). We baseline the MUA for each trial by average activity during 500 ms prior to first tone. We then normalize the recorded activity for each time point of a given trial for a given electrode by taking SEM across all trial recordings over that electrode and then average the normalized baseline activity across electrodes. For residual activity in particular, we consider 50 ms time window preceding each successive stimulus (from second tone to fifth tone) and average across all prestimulus windows. We exclude the first tone because residual activity is an after effect of preceding stimulus within short time interval. For simulated residual activity we simply average across all prestimulus windows over all trials and realizations of the model. To get the time collapsed residual activity we average the residual activity over the whole 50 msec time period to get the violin plots in Figure 5 c and d. The statistical significance reported in this section is tested by two-sample Kolmogorov–Smirnov test.

## Data Availability

Neurophysiological recordings of macaque that are used in this study for validation were presented in reference (14) and the data is available in Figshare (https://doi.org/10.6084/m9.figshare.26980630). The results from human fMRI study shown here are presented in reference (7) and corresponding data is also available in Figshare (https://doi.org/10.6084/m9.figshare.25927939)

## Code Availability

The model presented in this work is part of the Neuroblox computational neuroscience platform (code: https://neuroblox.ai/code ; documentation: https://neuroblox.ai/docs). An easy-to-use GUI, tutorials, and a standalone implementation for reproducing the results in this work are available at https://neuroblox.ai/pubs.

## Author Contributions Statement

Conceptualization: AP, HHS, LRMP, EKM and RG; Data curation: SLB, YX and EKM; Formal analysis: AP and RG; Funding acquisition: HHS, LRMP, EKM and RG; Investigation: AP and RG; Methodology: AP and RG; Project administration: LRMP; Software: AP, HO, MP and VD; Supervision: HHS, LRMP, EKM and RG; Validation: AP; Visualization: AP, LRMP and RG; Writing - original draft: AP and RG; Writing - review & editing: AP, SLB, HHS, LRMP, EKM and RG.

## Acknowledgements

We thank Yohan John for his valuable inputs in reviewing and editing of the manuscript. The research presented here was funded by the Baszucki Foundation, United States (LRMP). This work was supported in part by the Office of Naval Research, United States : N00014-21-1-2290 (RG). The experiments were funded by Office of Naval Research, United States : MURI N00014-23-1-2768 and MURI W911NF2410228, National Institute of Mental Health : 1R01MH131715-01, Freedom Together Foundation, and The Picower Institute for Learning and Memory (EKM).

## SUPPLEMENTARY MATERIAL

**Figure S1:**
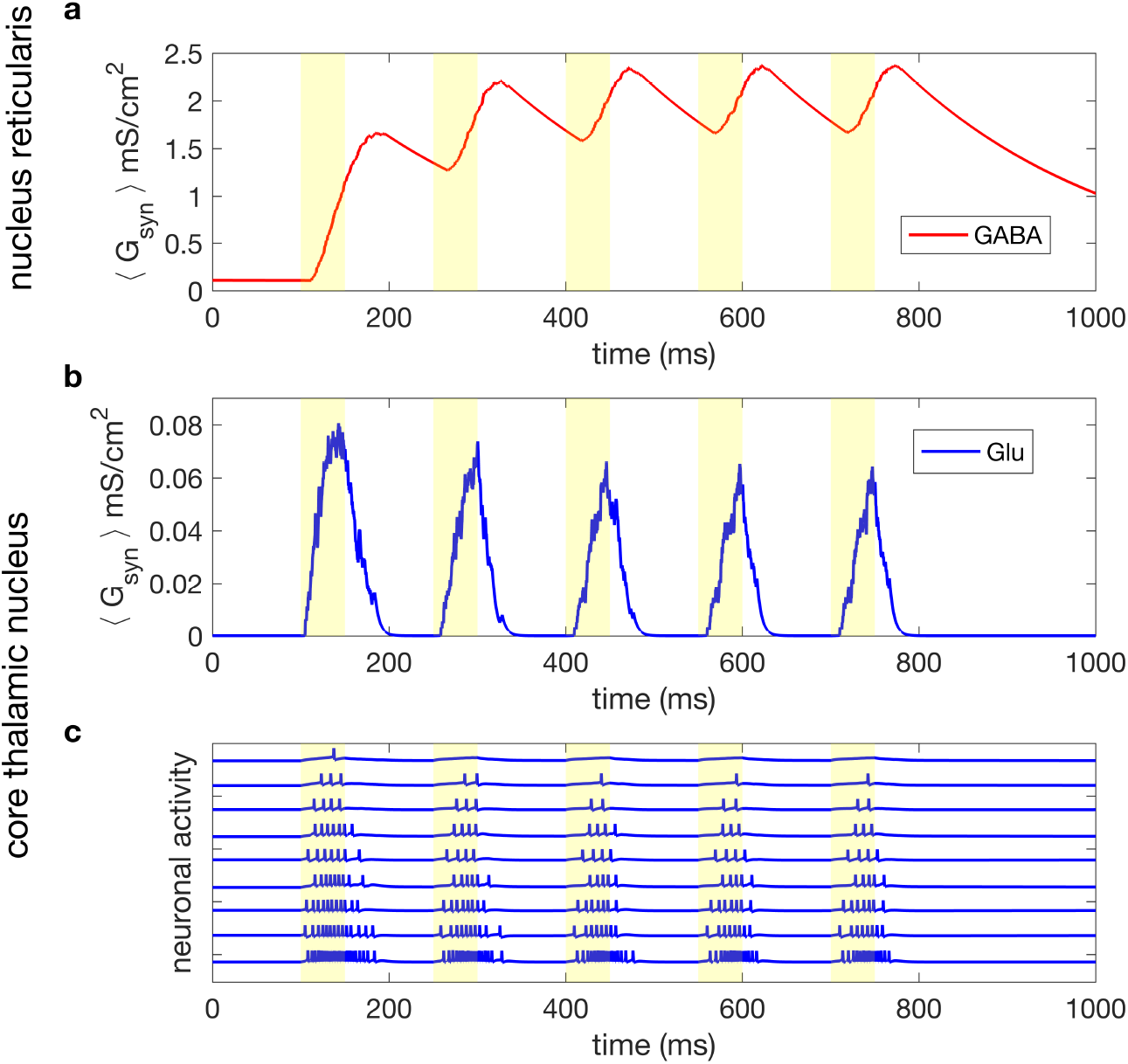
Nucleus Reticularis suppresses core thalamic activity response to repetitive stimulus. (a) Shows average synaptic conductance for GABAergic synapses between Nrt and core thalamic nucleus. Due to long time constants (300 ms) the inhibition builds up for successive auditory stimuli. (b) Simulated field potential in core thalamic nucleus showing attenuation in response to successive stimuli due to the inhibition build up. (c) Simulated spiking activity in core thalamic nucleus in response to the successive repetitive stimuli. The activity is progressively suppressed due to inhibition build up from nucleus reticularis shown in (a).

**Figure S2:**
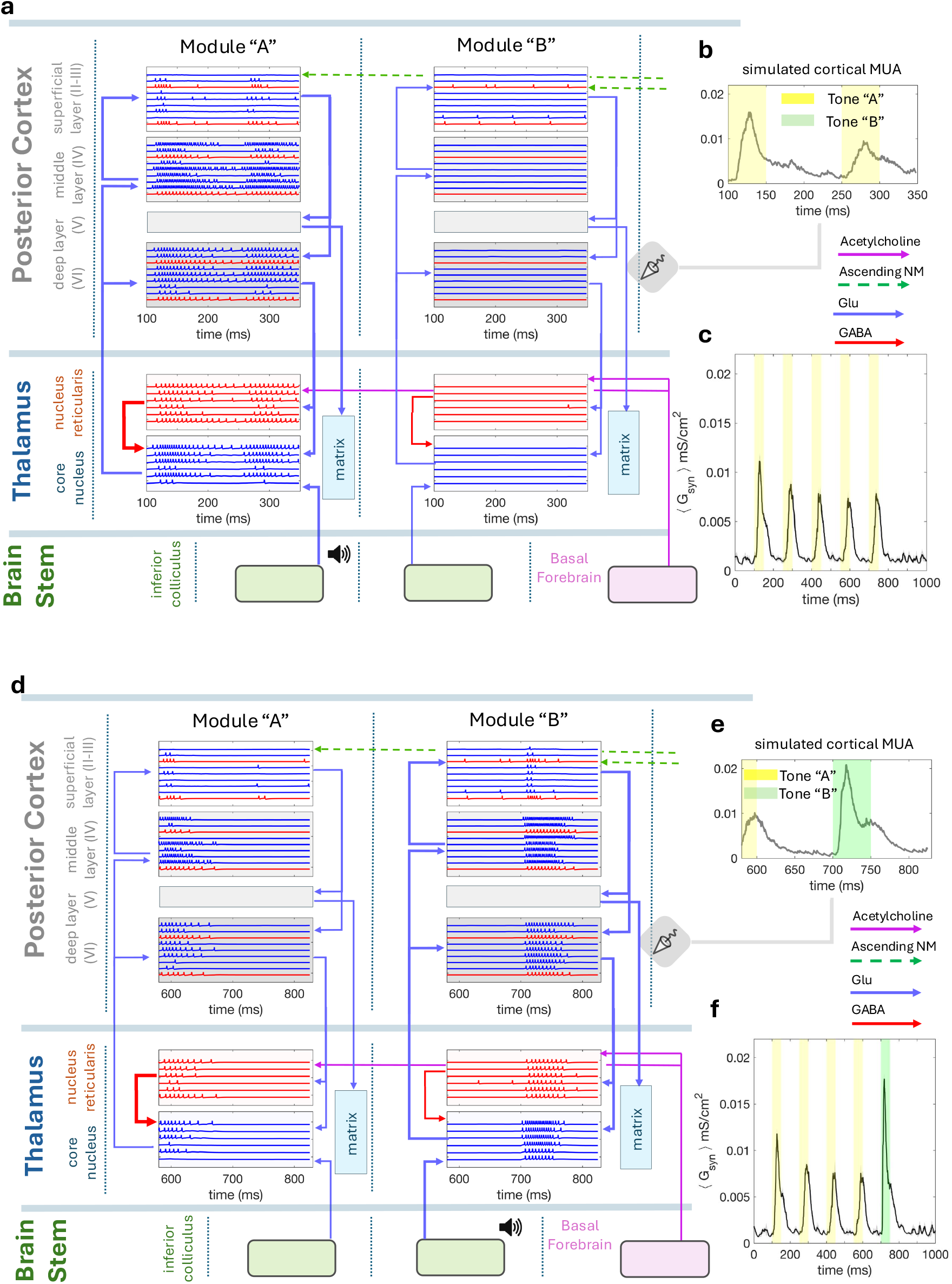
Simulation of a single oddball trial showing attenuation of response for regular tones (tone A) and uptick in response for oddball tone (tone B). We show simulation of two trial of odd ball task (5 successive tones) in awake state. Each tone is 50 ms and inter-tone interval is 100 ms. After the trial there is a long interval of 500 ms. (a-c) Shows response of the core thalamic circuit to a regular trial (AAAAA). (d-f) Shows response of the same circuit to a global oddball trial (AAAAB). (a) & (d) show spatio-temporal spiking activity pattern in thalamic nuclei and cortical layers (excluding matrix and deep cortical layer V as they are involved in the matrix thalamic loop, see Figure S5). (b) & (e) Show the simulated field potential response corresponding spiking activity from the cortical region. (c) & (f) Show the field potential response for the single auditory trial, regular and oddball respectively. Note the attenuation of amplitude for repetitive tones and an uptick for the odd tone. This circuit model focuses on the cortico-thalamic loops which might be majorly involved in passive sensory perception tasks, and associated brain stem regions (Inferior Colliculus, ASC1 (Pathak et al 2024), Basal Forebrain). Cortico-thalamic loops are of two kinds: core and matrix This figure shows the core part of the cortico-thalamic circuit involving the posterior cortex (primary auditory), as core thalamic circuit mainly involves primary sensory cortical regions. Core loop is topographically conserved across all cortical layers (superficial, middle, deep) and thalamic nuclei. Hence it is also called specific loop. Auditory stimulus enters through Inferior Colliculus. Different tones (frequencies) are mapped onto different anatomical modules from IC to core thalamus and to the cortical layers. After each tone, specific module gets activated (A or B). The nucleus reticularis in thalamus have long inhibitory time scales (300 ms) which keeps suppressing subsequent activities, thus the ‘downward step’ effect seen in the empirical data (see Figure 4, main text). The odd tone is not inhibited as it enters a separate module.

**Figure S3:**
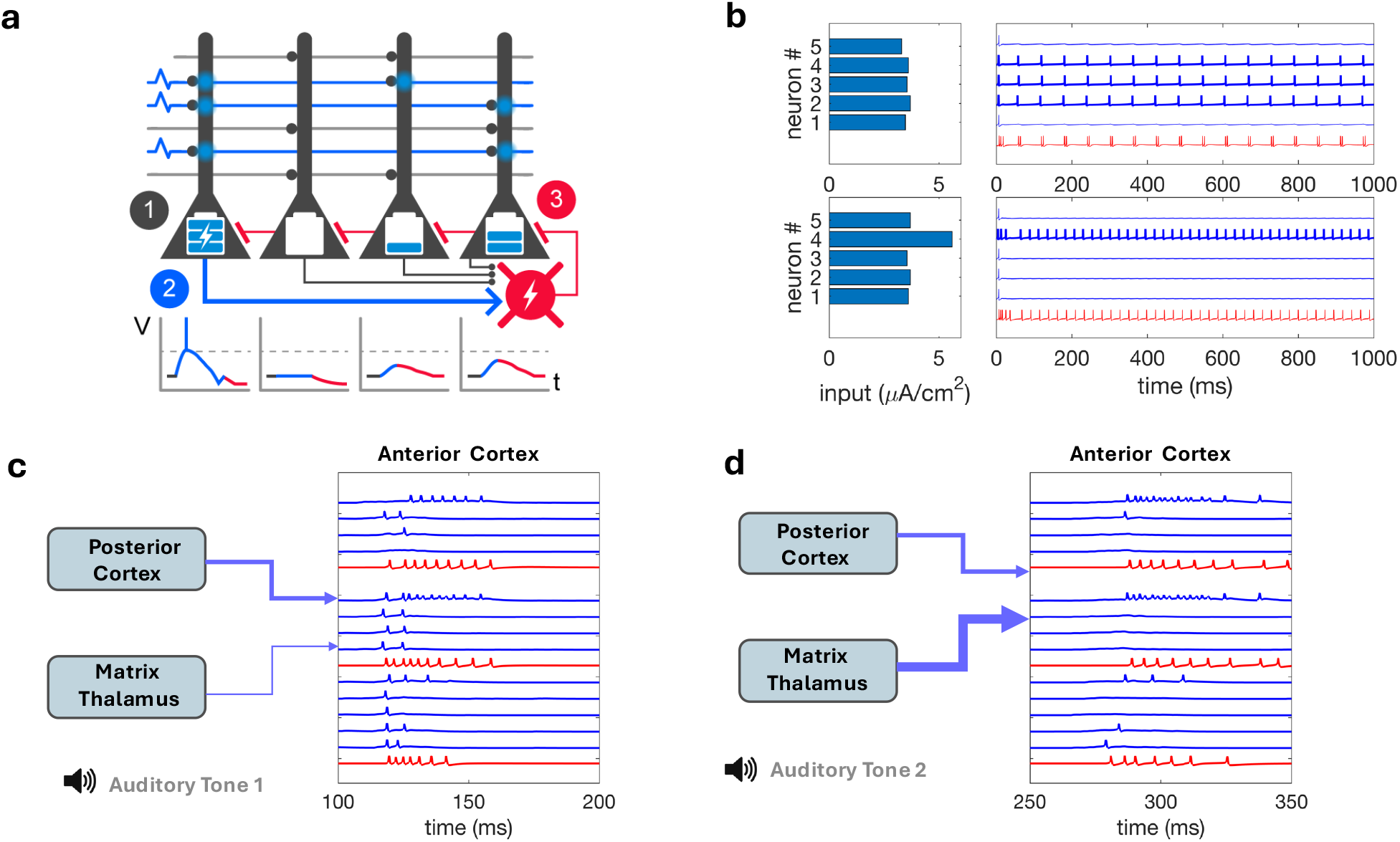
Matrix thalamic projections to cortex target not layer IV (unlike core thalamic projections), but layer I, which also receives input from cortico-cortical projections. Here the overlap between cortico-cortical and thalamo-cortical projections plays an important role. Target neurons in anterior cortex that get activated by cortical inputs from posterior cortex after first stimulus, get more susceptible to subsequent thalamic inputs during the subsequent stimuli due to short term augmentation (STA) seen in thalamo-cortical synapses [1, 2]. That causes a subset of neurons that get cortical input receive significantly higher inputs added by thalamic projections. This leads to inhibition of the remaining neurons through the feedback lateral inhibition seen in the LFLIC circuit [4]. (a) Shows a single LFLIC circuit that acts as a biomemetic computational primitive underlying the superficial cortical layer. (b) Demonstartes the response of a single LFLIC circuit neurons to an evenly distributed input (top) compared to a skewed input (bottom). Evenly distributed inputs cause more neurons to be active simultaneously while a skewed distribution leads to significantly sparser activity, even though the total input to the circuit is higher in the latter case. This mechanism plays out when matrix thalamus projections to anterior cortex overlap with the cortical projections, as illustrated in (c). Thalamic synapses into anterior cortex get augmented after the first stimulus, and for successive stimuli these augmented thalamic inputs reduce the active neuron population through lateral inhibition mechanism in the superficial cortex [4].

**Figure S4:**
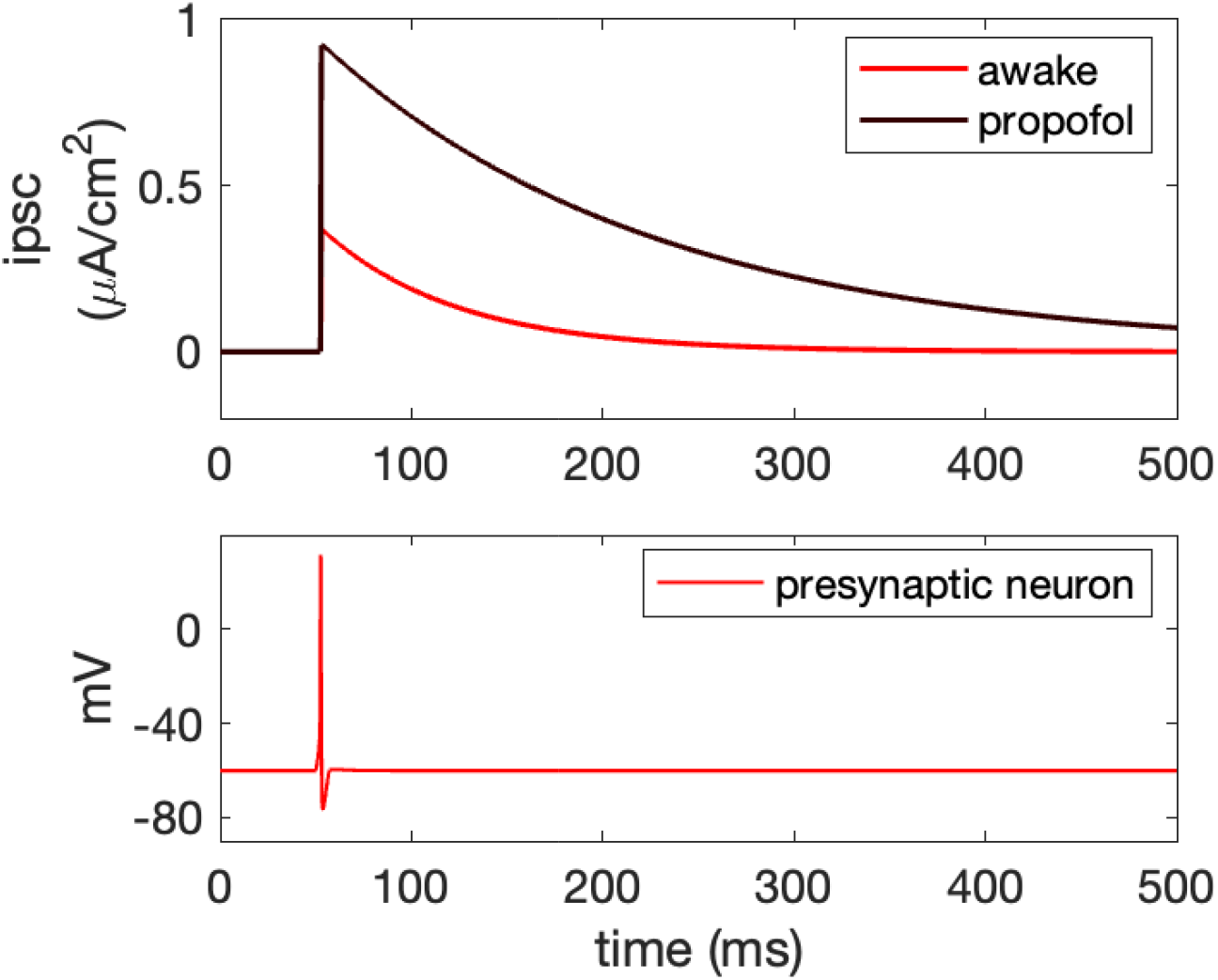
Inhibitory post synaptic currents in GABA-A receptors (top) caused by spiking of a GABAergic neuron (bottom). Propofol modulation causes the IPSCs get stronger in amplitude with an increase in the decay time.

**Figure S5:**
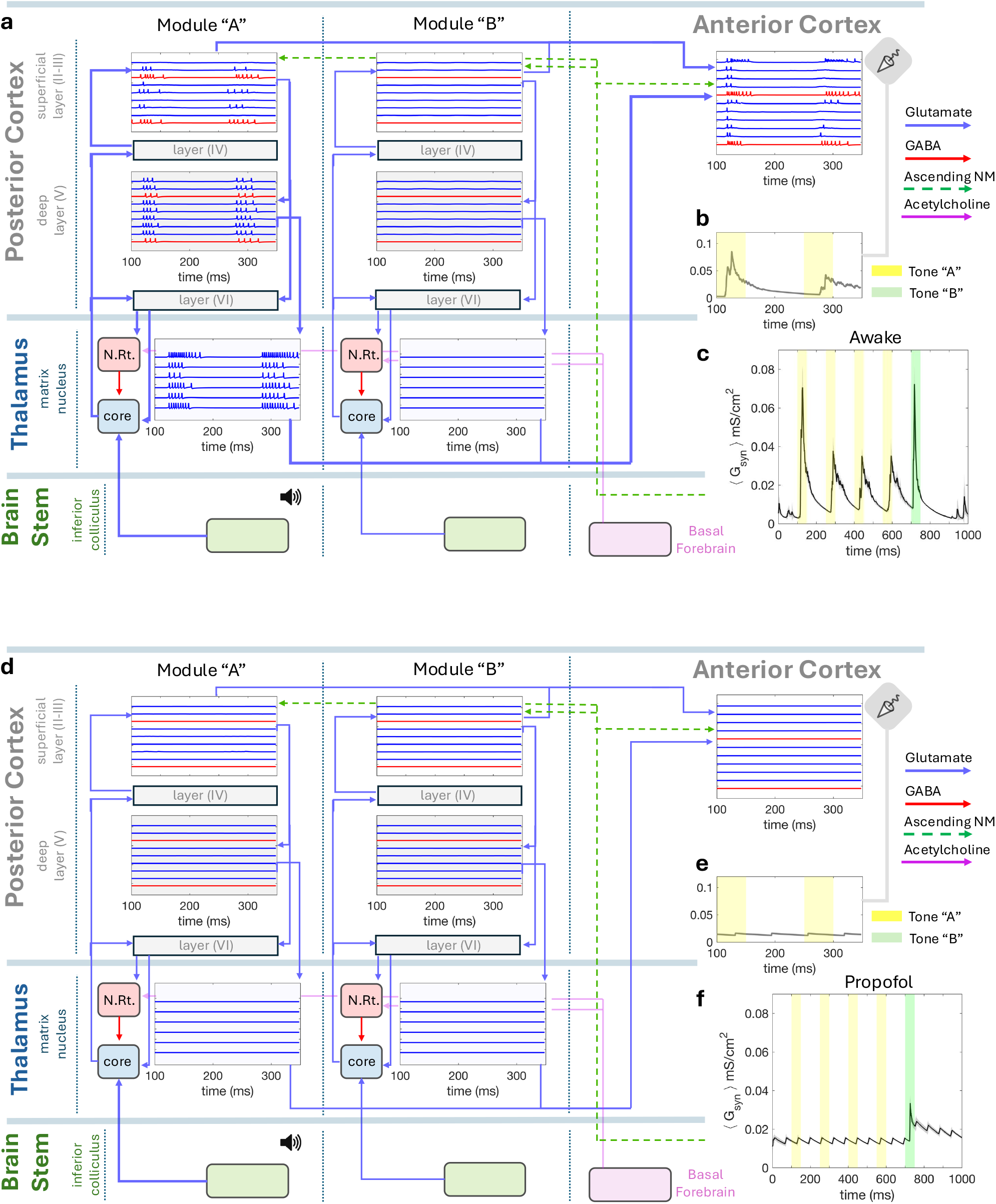
Spiking activity response in the matrix cortico-thalamic circuit during (a-c) awake and (d-f) anesthetized states. This complements the core thalamic ciruit shown in Figure 2, main text. In the awake state (A) we observe in the anterior cortex the lateral inhibitory effect from first to second stimuli, caused by overlapping inputs from posterior cortex and matrix thalamus (see Figure S3). In the anesthetized state (B), the superficial layer 2-3 of posterior cortex have almost completely suppressed activity. Since superficial layer 2-3 is the common connective component between the core circuit and the matrix circuit, suppression of superficial layer activity causes dissociation between the core and the matrix circuits [3]. Hence, the convergent input to anterior cortex are stopped in anesthetic state. Therefore we see no response in the anterior cortex for either regular or oddball stimuli, as opposed to posterior cortex where we see strong response for oddball in the anesthetic state (see Figure 4, main text

**Figure S6:**
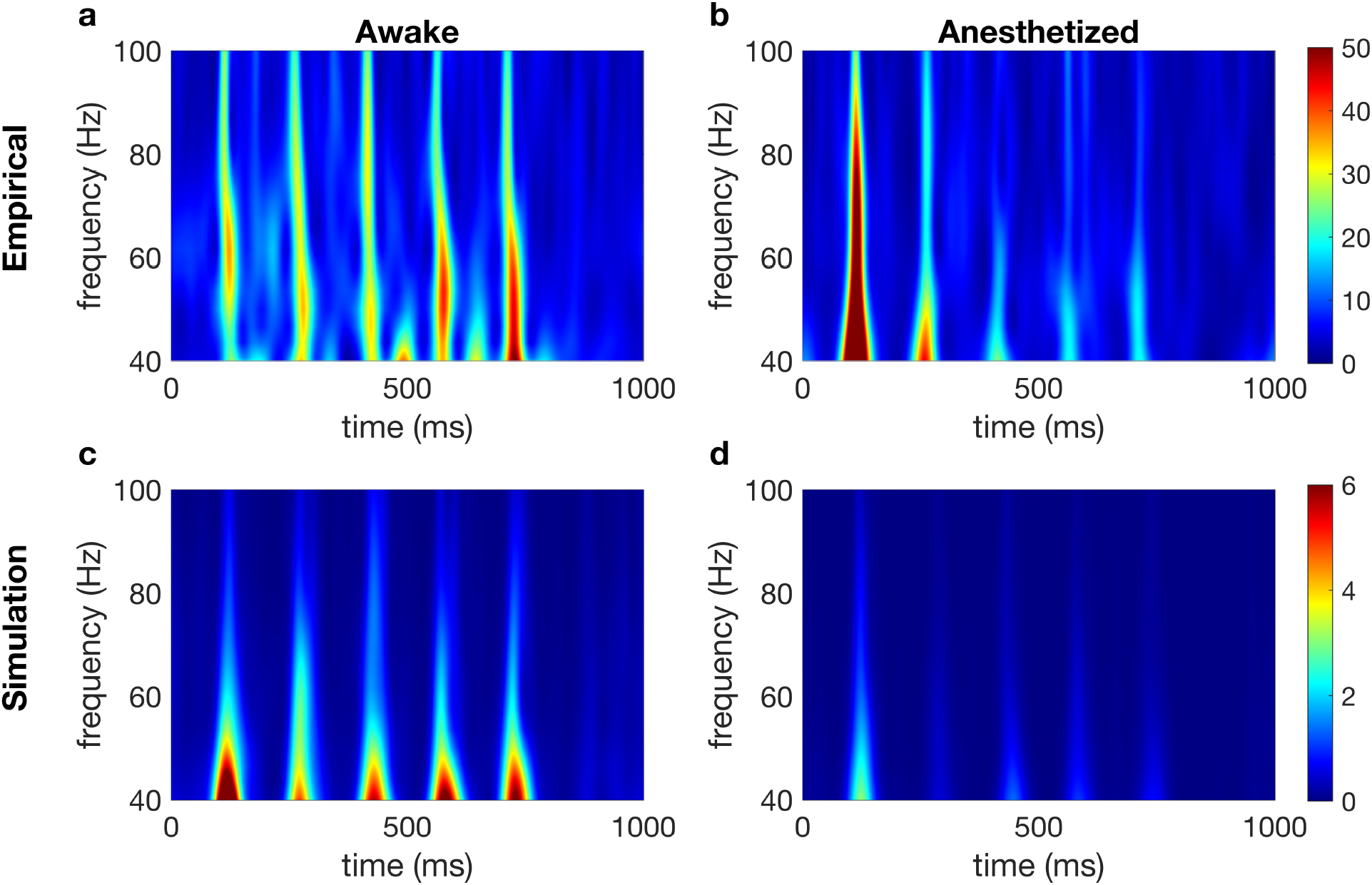
Suppression of gamma band in posterior cortex due to propofol induced anesthesia. Panels (a-d) show spectrogram for the empirical (a-b) and simulated (c-d) MUAs during successive repeated tones of the passive auditory task, averaged over all trials. Each burst of gamma band (40-100 Hz) is in response to a stimulus. Notice the suppression of gamma activity caused by propofol (b and d) as compared to awake state (a and c).

**Figure S7:**
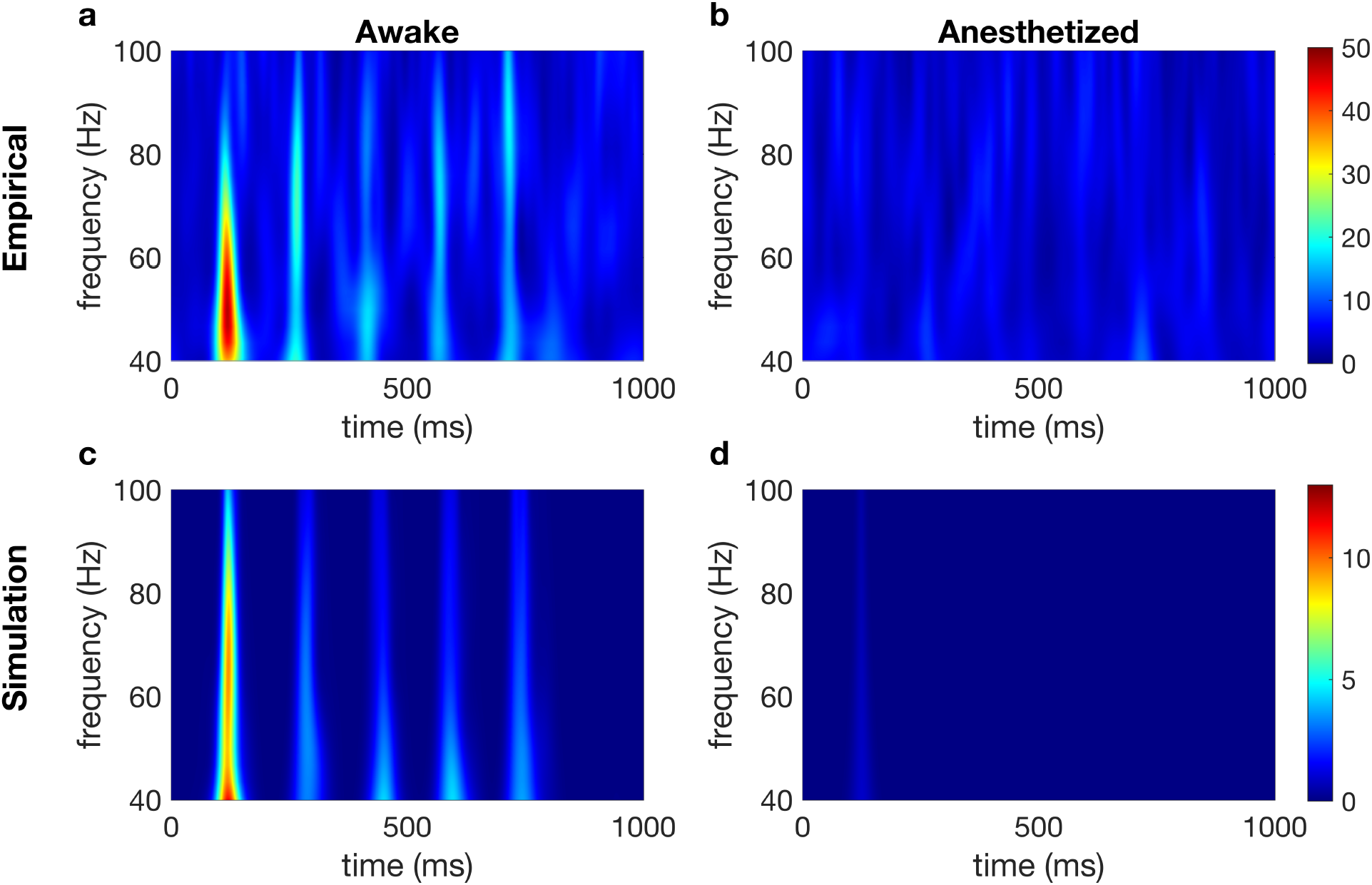
Suppression of gamma band in anterior cortex due to propofol induced anesthesia. Panels (a-d) show spectrogram for the empirical (a-b) and simulated (c-d) MUAs during successive repeated tones of the passive auditory task, averaged over all trials. Each burst of gamma band (40-100 Hz) is in response to a stimulus. Notice the suppression of gamma activity caused by propofol (b and d) as compared to awake state (a and c).

**Figure S8:**
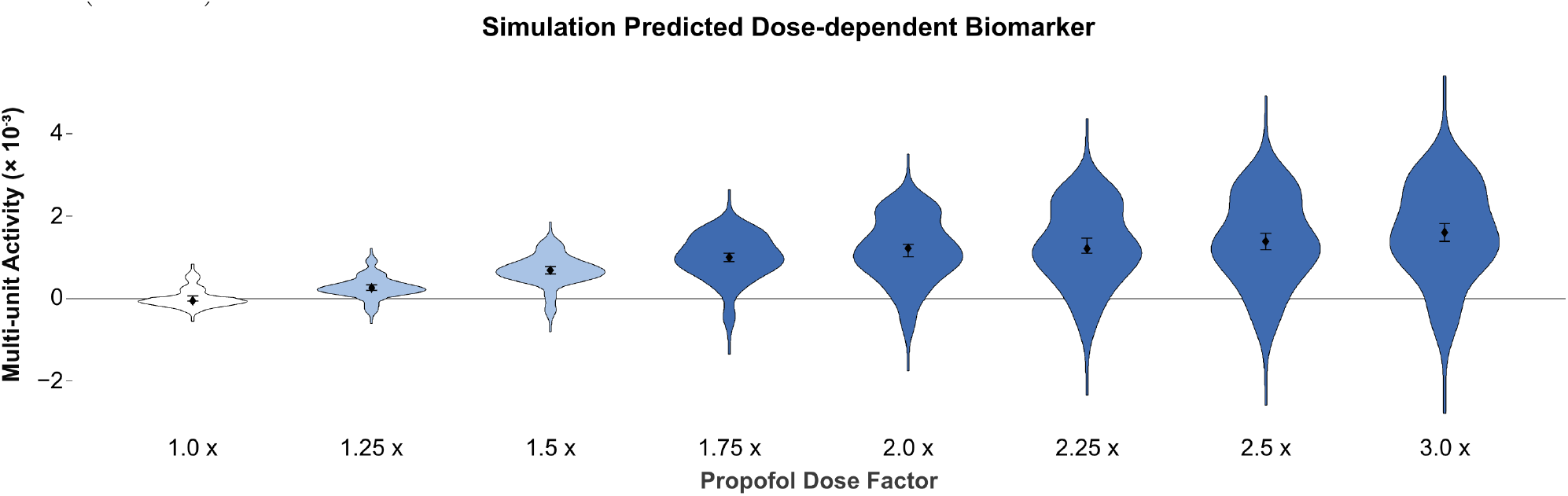
Suppression of gamma band in anterior cortex due to propofol induced anesthesia. Panels (a-d) show spectrogram for the empirical (a-b) and simulated (c-d) MUAs during successive repeated tones of the passive auditory task, averaged over all trials. Each burst of gamma band (40-100 Hz) is in response to a stimulus. Notice the suppression of gamma activity caused by propofol (b and d) as compared to awake state (a and c).

### Robustness Perturbation Tests

Following are the results of robustness perturbation tests. The model is strongly perturbed with the key parameters such as connection densities, synaptic weights and background inputs to each of the components are simultaneously changed ranging from 20% to 50%. These results show that the model sustains the primary behavior in the control and anesthetized states.

**Figure S9:**
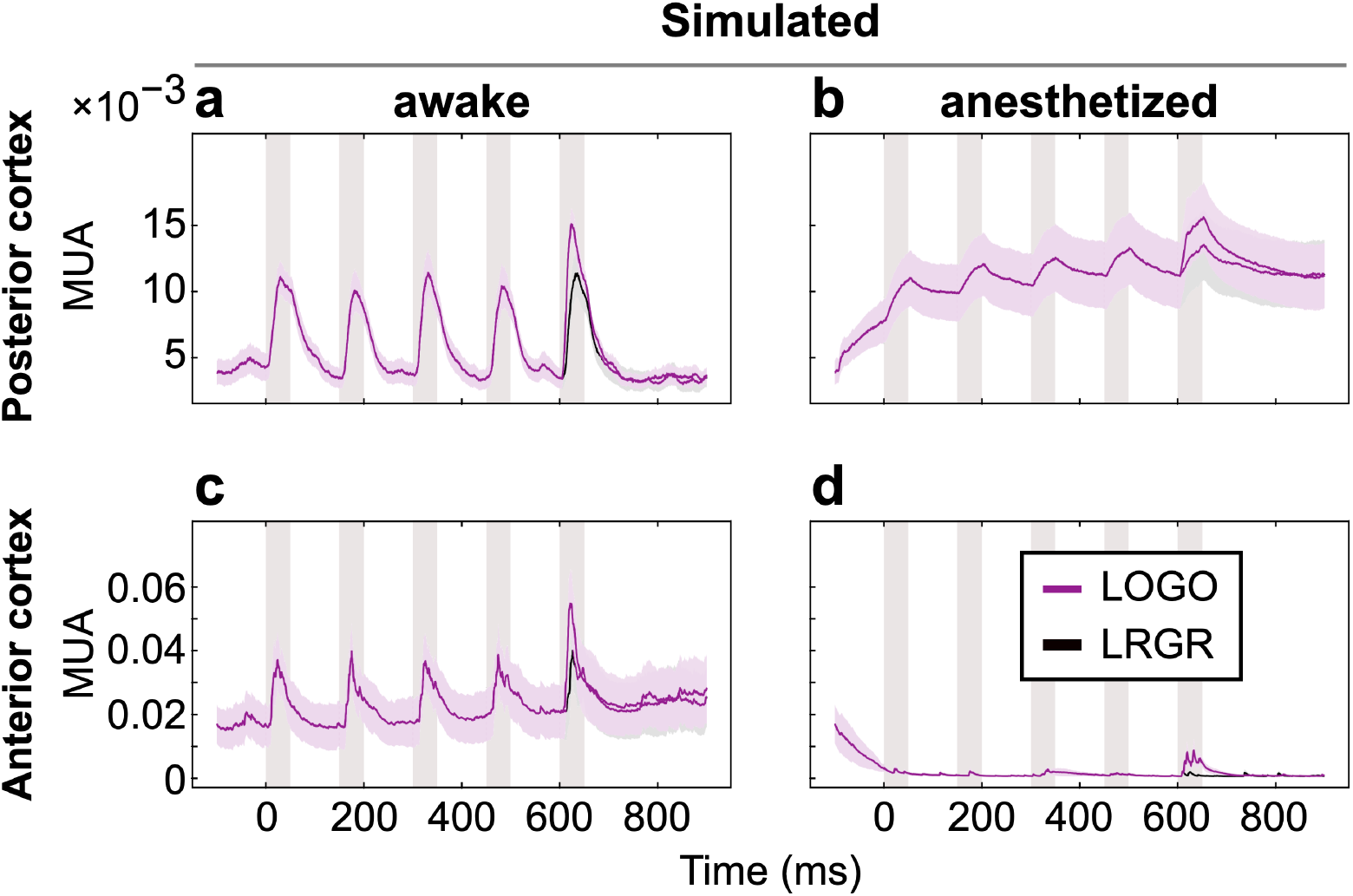
Perturbation of model retains the electrophysiological responses to stimulation. Perturbed model simulation responses to tone sequences in both posterior (a-b) and anterior (c-d) cortical regions, under control awake and anesthetized conditions (see Figure 3 in Main Text). Shaded pink regions indicate duration of each auditory stimulus. The simulation was not trained on this or any macaque data. Instead, the bottom-up simulation was presented with the same auditory tone sequence stimuli that were presented to the macaques (see Fig 2), and the resulting simulated physiology was produced.

**Figure S10:**
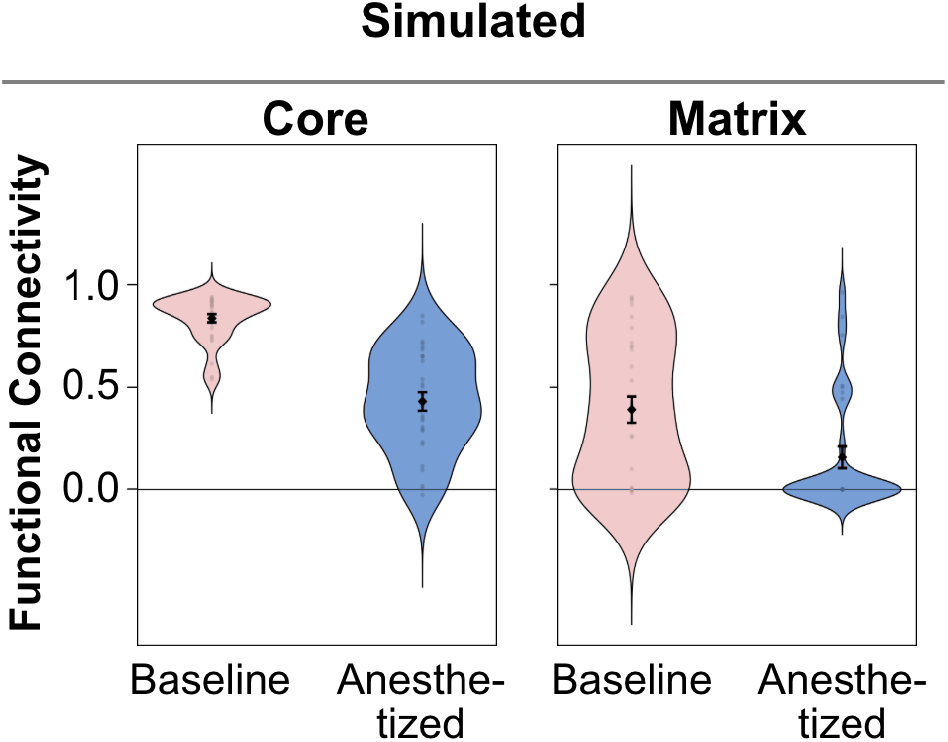
Perturbed model retains the collapse of matrix loop functional connectivity under anesthesia. Simulated functional connectivities of core and matrix loops shown from perturbed model (see Figure 4 in the main text for comparison).

**Figure S11:**
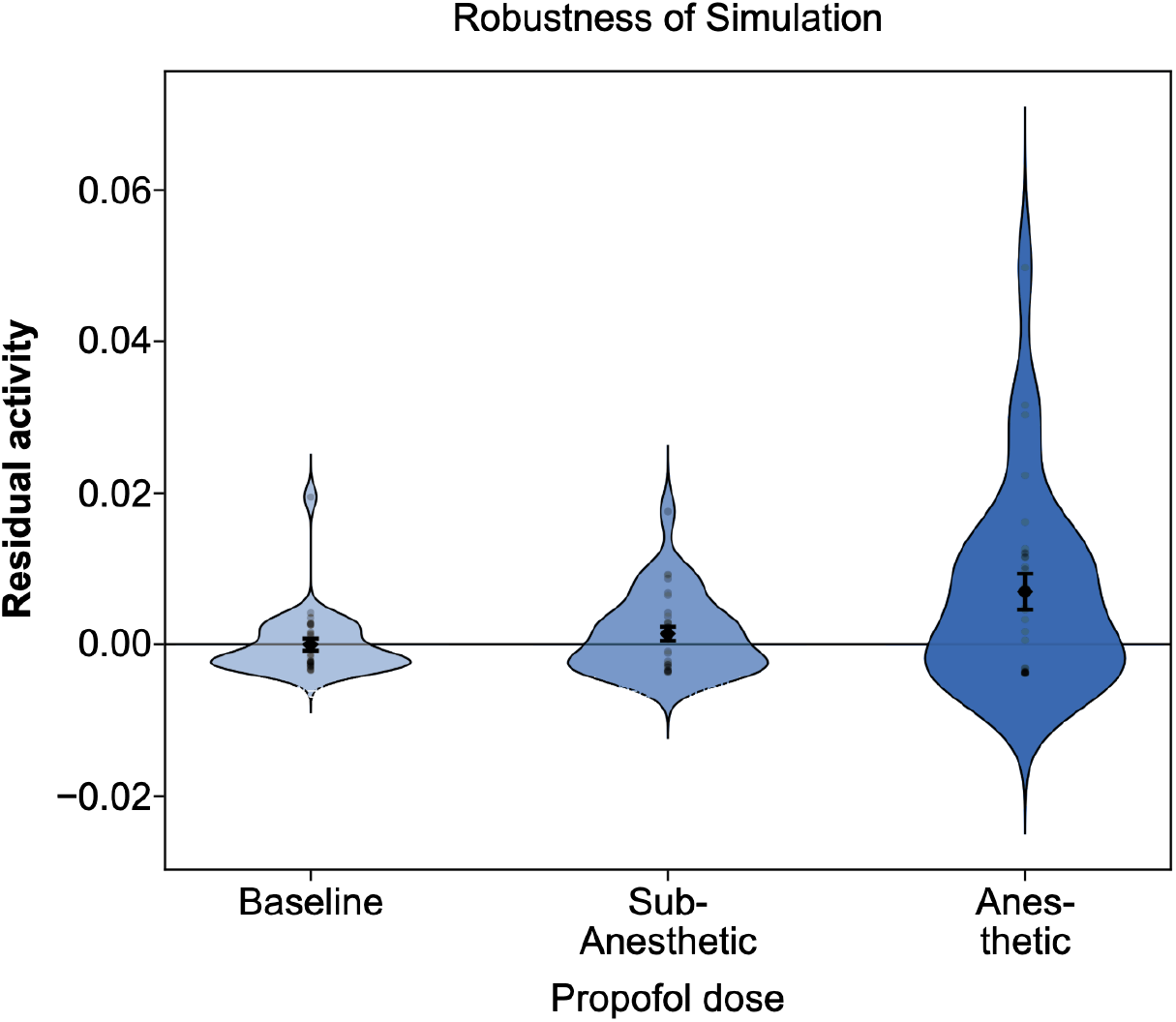
Perturbed model retains the residual activity that increases with anesthetic dose. Simulated residual activities in posterior cortex from perturbed model (see Figure 5 in the main text for comparison).

