## Supplementary Information for "Mechanistic simulation identifies predictive dose-dependent biomarkers of propofol anesthesia"

### SUPPLEMENTARY MATERIAL

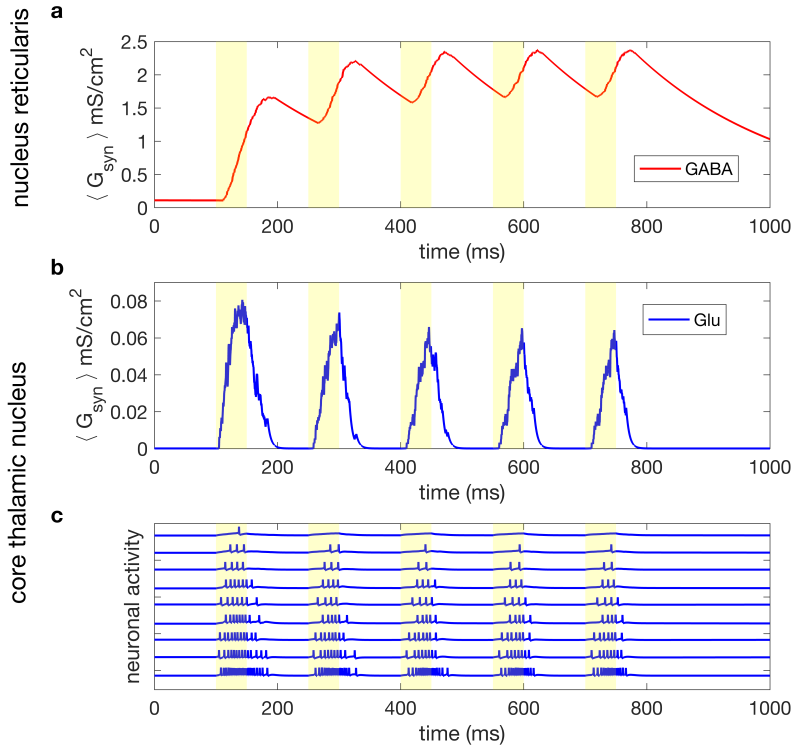


Figure S1: Nucleus Reticularis suppresses core thalamic activity response to repetitive stimulus. (a) Shows average synaptic conductance for GABAergic synapses between Nrt and core thalamic nucleus. Due to long time constants ( 300 ms) the inhibition builds up for successive auditory stimuli. (b) Simulated field potential in core thalamic nucleus showing attenuation in response to successive stimuli due to the inhibition build up. (c) Simulated spiking activity in core thalamic nucleus in response to the successive repetitive stimuli. The activity is progressively suppressed due to inhibition build up from nucleus reticularis shown in (a).

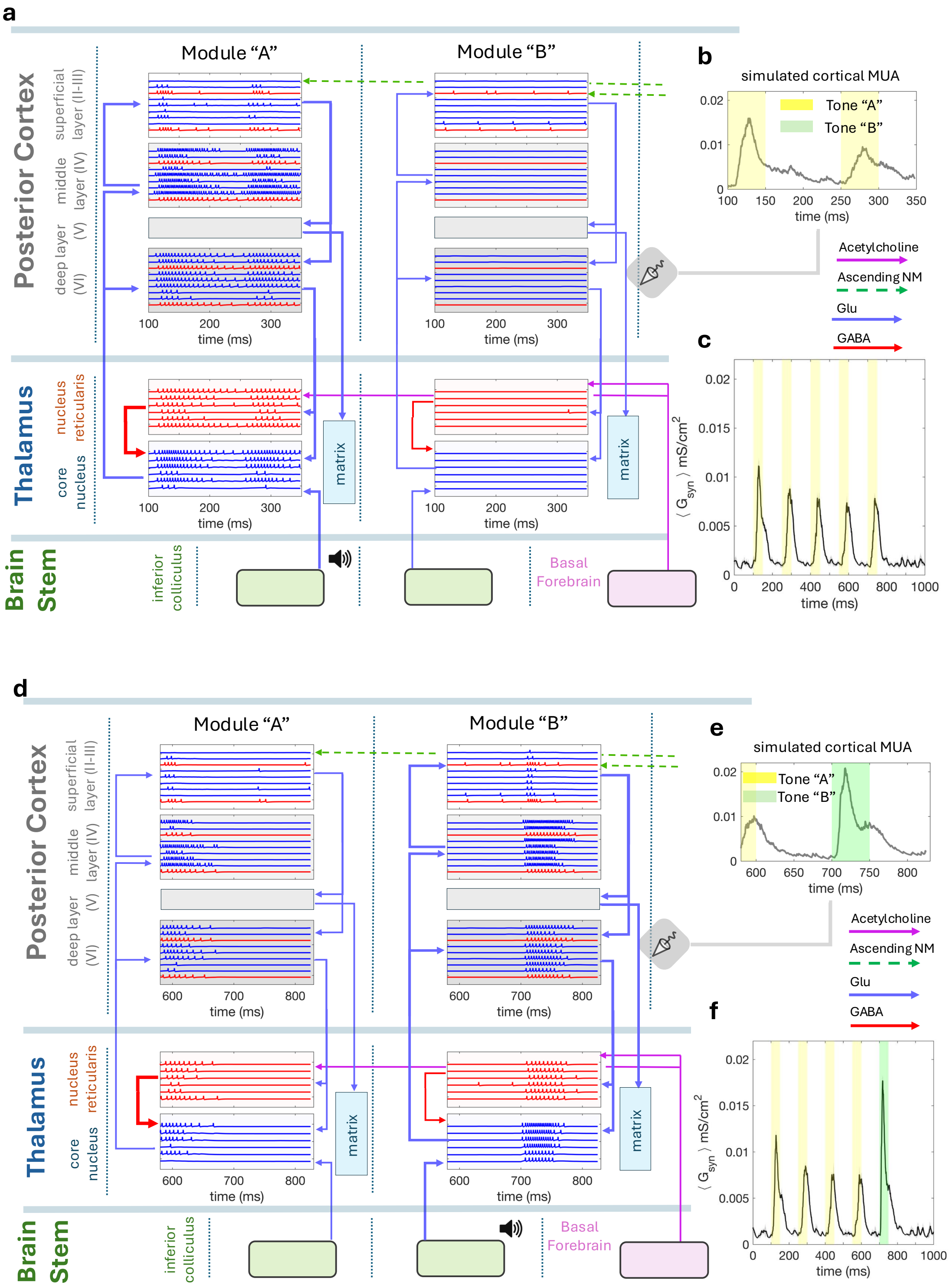
a
Module "A"
Module "B"
Posterior Cortex
superficial layer (II-III)
middle layer (IV)
deep layer (V)
(VI)
time (ms)
Thalamus
nucleus reticularis
core nucleus
matrix
Brain Stem
inferior colliculus
Basal Forebrain
b
simulated cortical MUA
Tone "A"
Tone "B"
Acetylcholine
Ascending NM
Glu
GABA
c
$\langle G_{syn} \rangle$ mS/cm$^2$
d
e
f

Figure S2

Figure S2: Simulation of a single oddball trial showing attenuation of response for regular tones (tone A) and uptick in response for oddball tone (tone B). We show simulation of two trial of odd ball task (5 successive tones) in awake state. Each tone is 50 ms and inter-tone interval is 100 ms. After the trial there is a long interval of 500 ms. (a-c) Shows response of the core thalamic circuit to a regular trial (AAAAA). (d-f) Shows response of the same circuit to a global oddball trial (AAAAB). (a) & (d) show spatio-temporal spiking activity pattern in thalamic nuclei and cortical layers (excluding matrix and deep cortical layer V as they are involved in the matrix thalamic loop, see Figure S5). (b) & (e) Show the simulated field potential response corresponding spiking activity from the cortical region. (c) & (f) Show the field potential response for the single auditory trial, regular and oddball respectively. Note the attenuation of amplitude for repetitive tones and an uptick for the odd tone. This circuit model focuses on the cortico-thalamic loops which might be majorly involved in passive sensory perception tasks, and associated brain stem regions (Inferior Colliculus, ASC1 (Pathak et al 2024), Basal Forebrain). Cortico-thalamic loops are of two kinds: core and matrix This figure shows the core part of the cortico-thalamic circuit involving the posterior cortex (primary auditory), as core thalamic circuit mainly involves primary sensory cortical regions. Core loop is topographically conserved across all cortical layers (superficial, middle, deep) and thalamic nuclei. Hence it is also called specific loop. Auditory stimulus enters through Inferior Colliculus. Different tones (frequencies) are mapped onto different anatomical modules from IC to core thalamus and to the cortical layers. After each tone, specific module gets activated (A or B). The nucleus reticularis in thalamus have long inhibitory time scales ( 300 ms) which keeps suppressing subsequent activities, thus the 'downward step' effect seen in the empirical data (see Figure 4, main text). The odd tone is not inhibited as it enters a separate module.

.

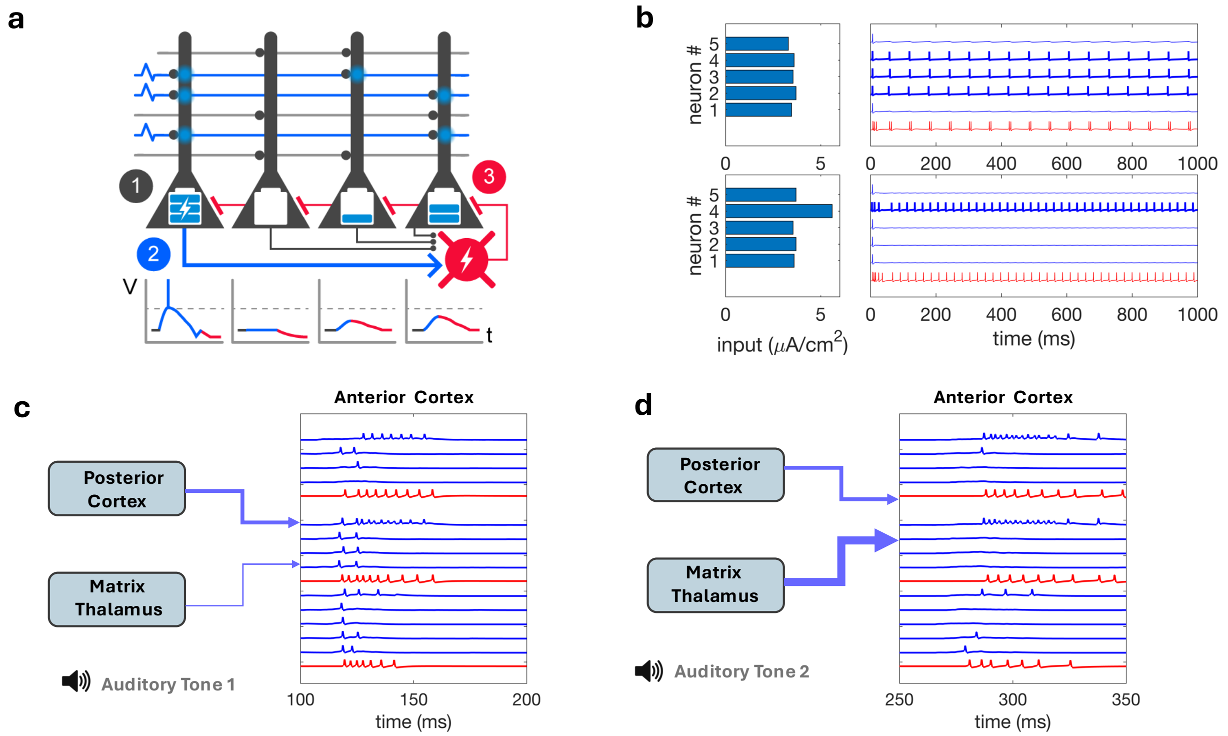


Figure S3: Matrix thalamic projections to cortex target not layer IV (unlike core thalamic projections), but layer I, which also receives input from cortico-cortical projections. Here the overlap between cortico-cortical and thalamo-cortical projections plays an important role. Target neurons in anterior cortex that get activated by cortical inputs from posterior cortex after first stimulus, get more susceptible to subsequent thalamic inputs during the subsequent stimuli due to short term augmentation (STA) seen in thalamo-cortical synapses [1, 2]. That causes a subset of neurons that get cortical input receive significantly higher inputs added by thalamic projections. This leads to inhibition of the remaining neurons through the feedback lateral inhibition seen in the LFLIC circuit [4]. (a) Shows a single LFLIC circuit that acts as a biomemetic computational primitive underlying the superficial cortical layer. (b) Demonstartes the response of a single LFLIC circuit neurons to an evenly distributed input (top) compared to a skewed input (bottom). Evenly distributed inputs cause more neurons to be active simultaneously while a skewed distribution leads to significantly sparser activity, even though the total input to the circuit is higher in the latter case. This mechanism plays out when matrix thalamus projections to anterior cortex overlap with the cortical projections, as illustrated in (c). Thalamic synapses into anterior cortex get augmented after the first stimulus, and for successive stimuli these augmented thalamic inputs reduce the active neuron population through lateral inhibition mechanism in the superficial cortex [4].

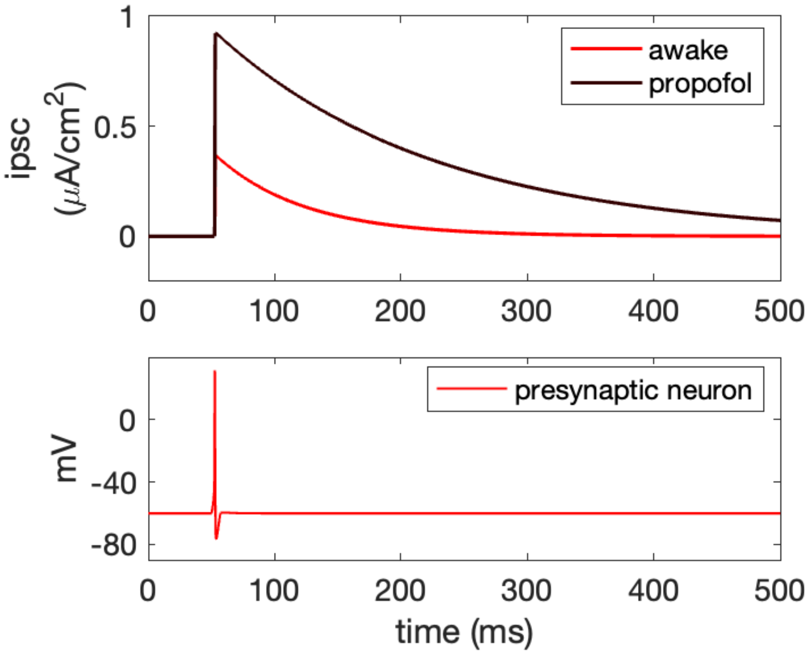


Figure S4: Inhibitory post synaptic currents in GABA-A receptors (top) caused by spiking of a GABAergic neuron (bottom). Propofol modulation causes the IPSCs get stronger in amplitude with an increase in the decay time.

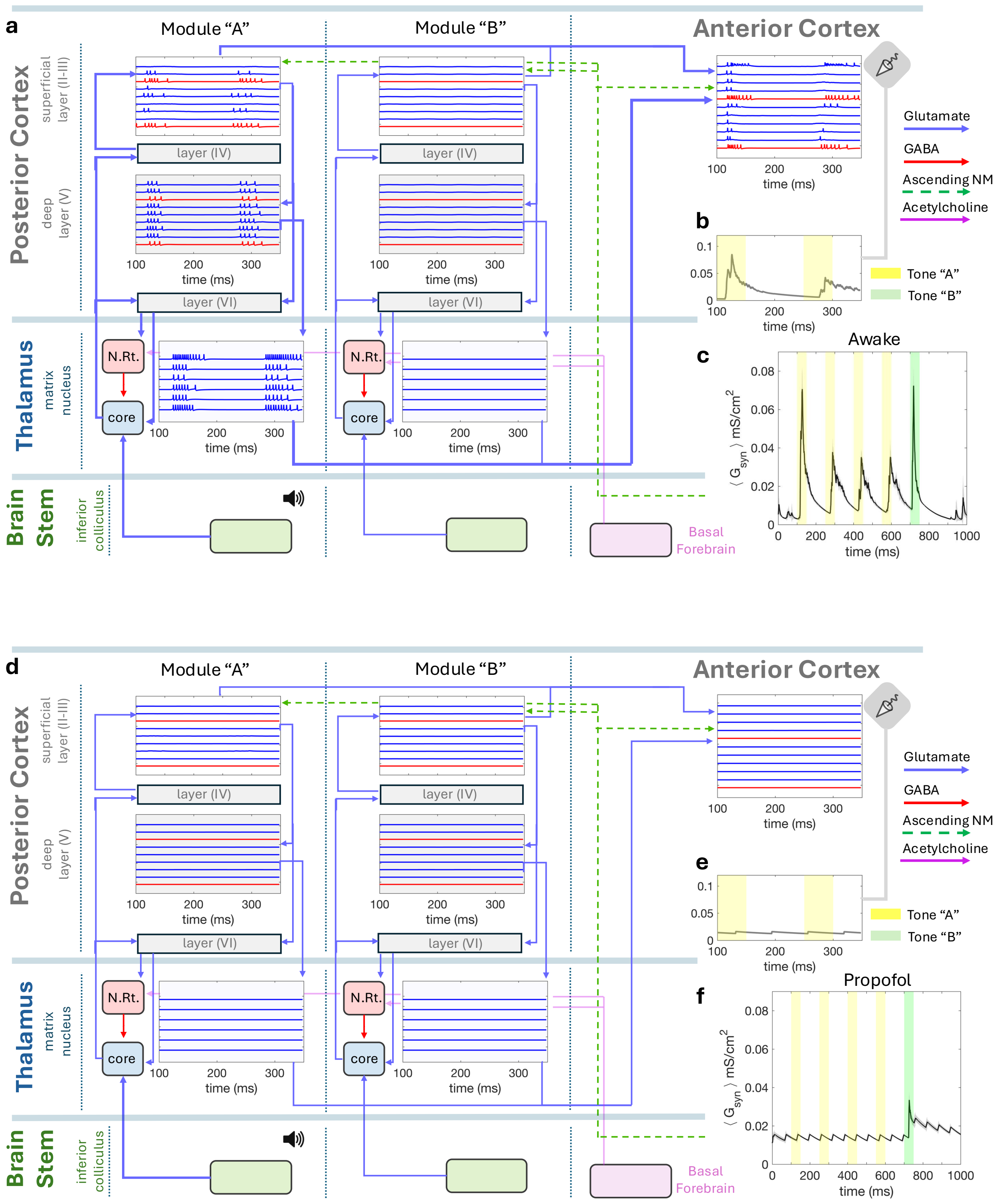

a
Module "A"
Module "B"
Anterior Cortex
Posterior Cortex
superficial layer (II-III)
deep layer (V)
layer (IV)
layer (VI)
time (ms)
Glutamate
GABA
Ascending NM
Acetylcholine
b
Tone "A"
Tone "B"
Awake
c
$\langle G_{syn} \rangle$ mS/cm$^2$
Thalamus
matrix nucleus
N.Rt.
core
Brain Stem
inferior colliculus
Basal Forebrain
d
e
Propofol
f


Figure S5

Figure S5: Spiking activity response in the matrix cortico-thalamic circuit during (a-c) awake and (d-f) anesthetized states. This complements the core thalamic ciruit shown in Figure 2, main text. In the awake state (A) we observe in the anterior cortex the lateral inhibitory effect from first to second stimuli, caused by overlapping inputs from posterior cortex and matrix thalamus (see Figure S3). In the anesthetized state (B), the superficial layer 2-3 of posterior cortex have almost completely suppressed activity. Since superficial layer 2-3 is the common connective component between the core circuit and the matrix circuit, suppression of superficial layer activity causes dissociation between the core and the matrix circuits [3]. Hence, the convergent input to anterior cortex are stopped in anesthetic state. Therefore we see no response in the anterior cortex for either regular or oddball stimuli, as opposed to posterior cortex where we see strong response for oddball in the anesthetic state (see Figure 4, main text

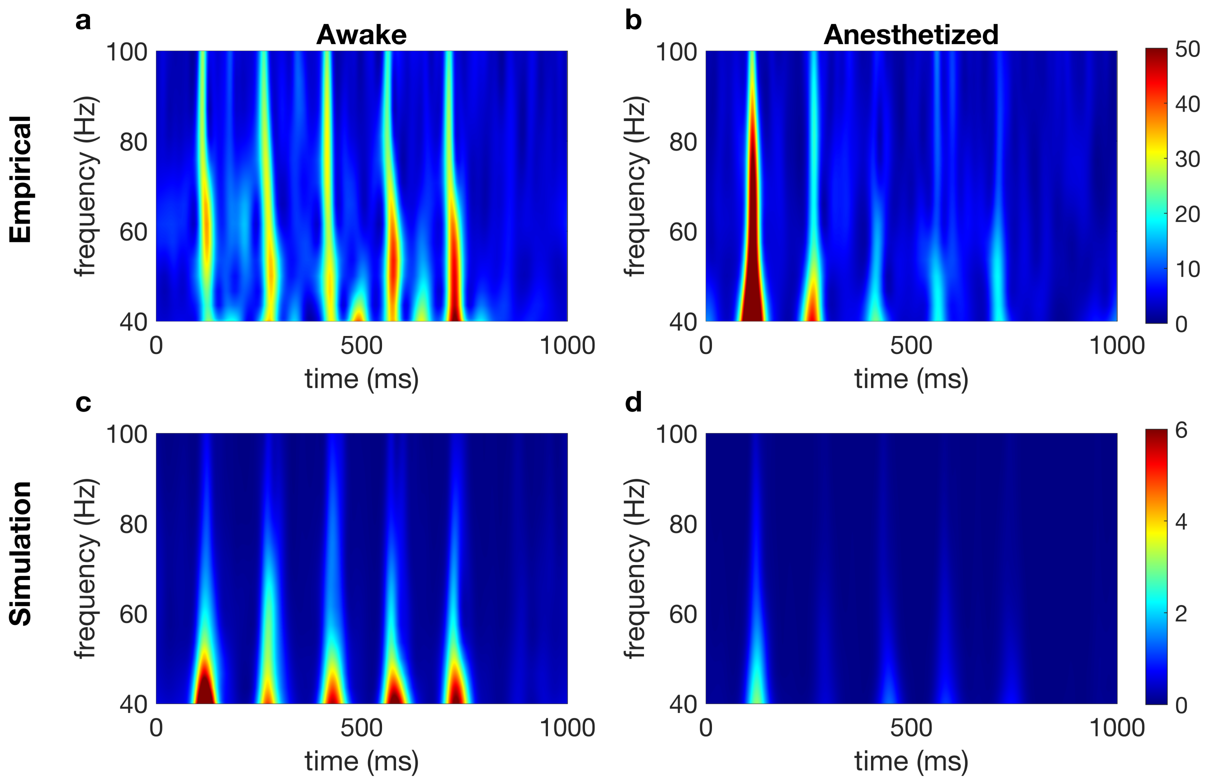


Figure S6: Suppression of gamma band in posterior cortex due to propofol induced anesthesia. Panels (a-d) show spectrogram for the empirical (a-b) and simulated (c-d) MUAs during successive repeated tones of the passive auditory task, averaged over all trials. Each burst of gamma band (40-100 Hz) is in response to a stimulus. Notice the suppression of gamma activity caused by propofol (b and d) as compared to awake state (a and c).

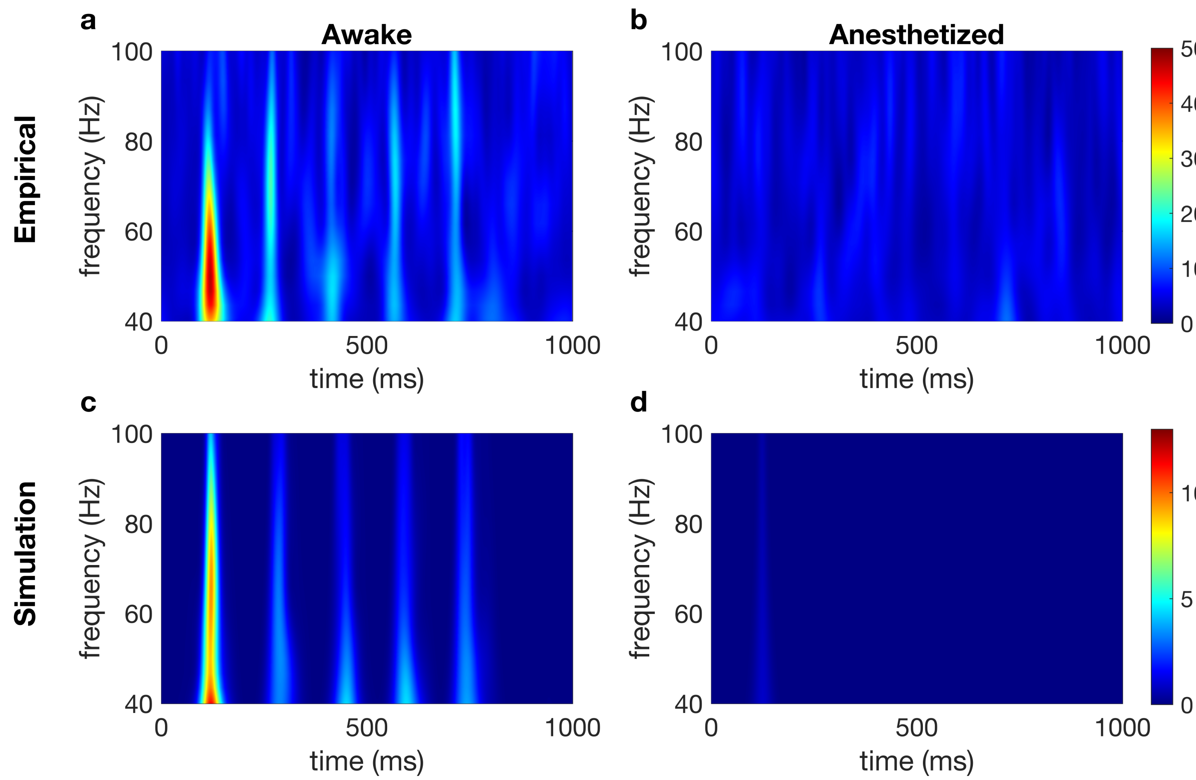


Figure S7: Suppression of gamma band in anterior cortex due to propofol induced anesthesia. Panels (a-d) show spectrogram for the empirical (a-b) and simulated (c-d) MUAs during successive repeated tones of the passive auditory task, averaged over all trials. Each burst of gamma band (40-100 Hz) is in response to a stimulus. Notice the suppression of gamma activity caused by propofol (b and d) as compared to awake state (a and c).

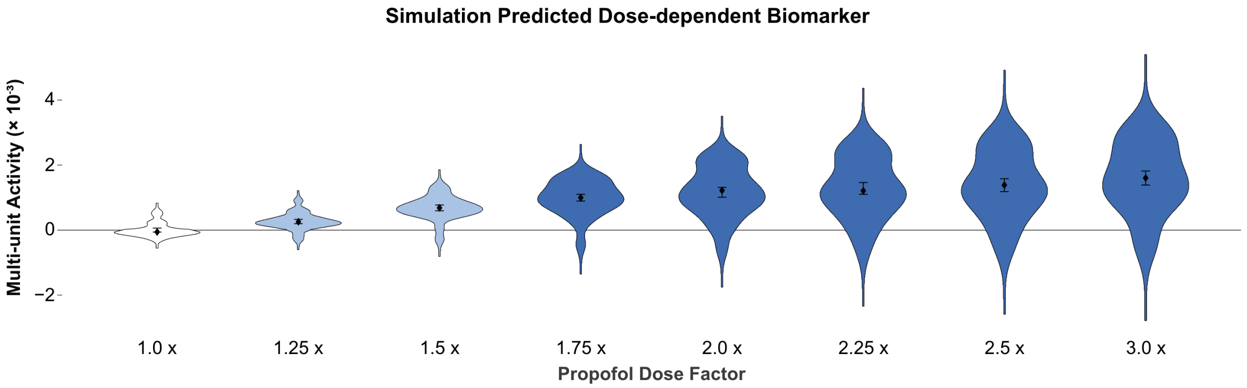


Figure S8: Suppression of gamma band in anterior cortex due to propofol induced anesthesia. Panels (a-d) show spectrogram for the empirical (a-b) and simulated (c-d) MUAs during successive repeated tones of the passive auditory task, averaged over all trials. Each burst of gamma band (40-100 Hz) is in response to a stimulus. Notice the suppression of gamma activity caused by propofol (b and d) as compared to awake state (a and c).

### Robustness Perturbation Tests

Following are the results of robustness perturbation tests. The model is strongly perturbed with the key parameters such as connection densities, synaptic weights and background inputs to each of the components are simultaneously changed ranging from 20% to 50%. These results show that the model sustains the primary behavior in the control and anesthetized states.

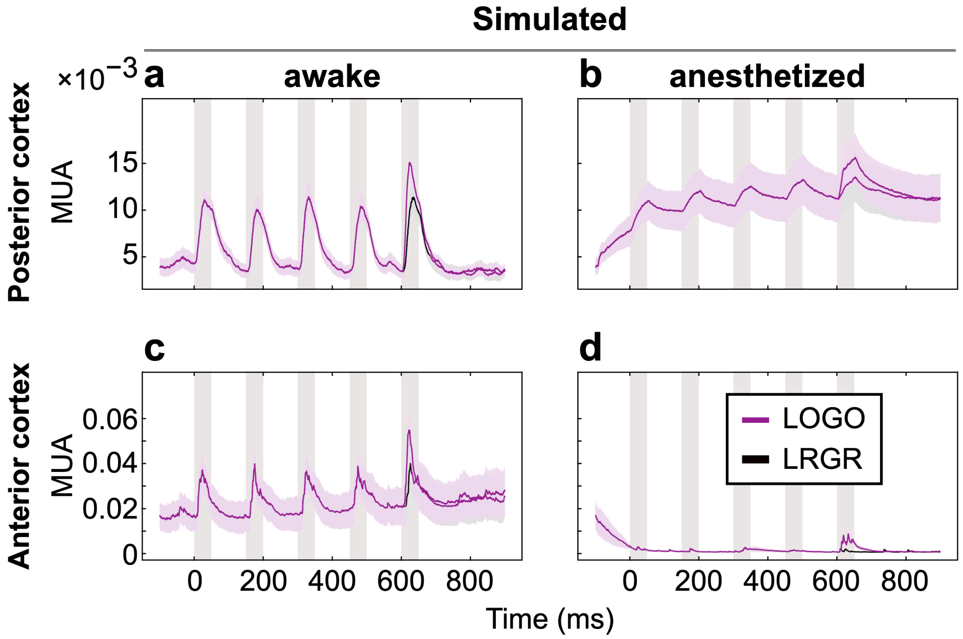


Figure S9: **Perturbation of model retains the electrophysiological responses to stimulation.** Perturbed model simulation responses to tone sequences in both posterior (a-b) and anterior (c-d) cortical regions, under control awake and anesthetized conditions (see Figure 3 in Main Text). Shaded pink regions indicate duration of each auditory stimulus. The simulation was not trained on this or any macaque data. Instead, the bottom-up simulation was presented with the same auditory tone sequence stimuli that were presented to the macaques (see Fig 2), and the resulting simulated physiology was produced.

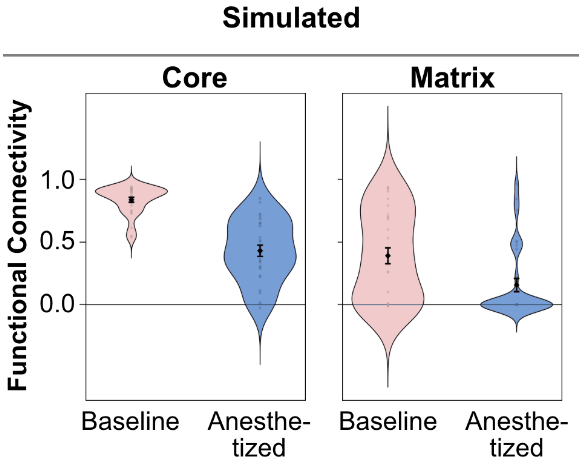


Figure S10: **Perturbed model retains the collapse of matrix loop functional connectivity under anesthesia**. Simulated functional connectivities of core and matrix loops shown from perturbed model (see Figure 4 in the main text for comparison).

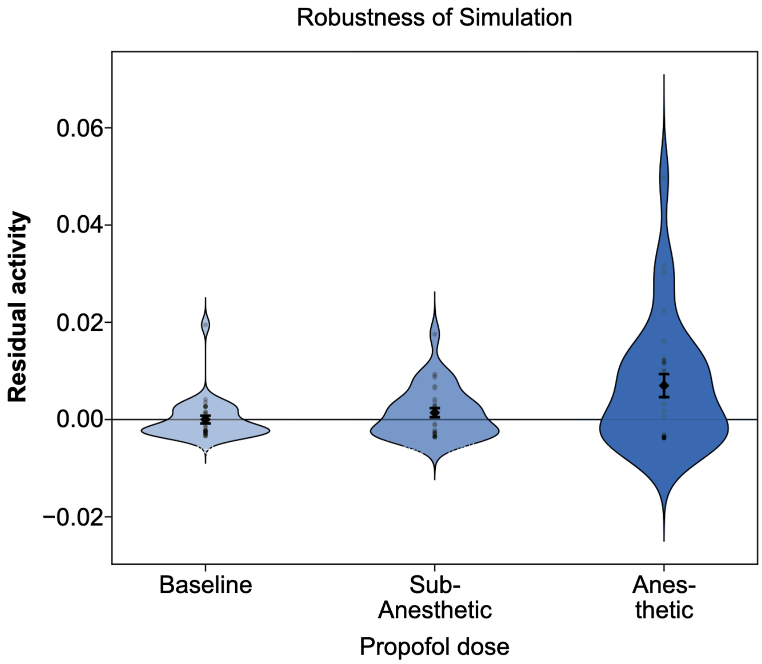


Figure S11: **Perturbed model retains the residual activity that increases with anesthetic dose**. Simulated residual activities in posterior cortex from perturbed model (see Figure 5 in the main text for comparison).
